# E-cadherin endocytosis is modulated by p120-catenin through the opposing actions of RhoA and Arf1

**DOI:** 10.1101/477760

**Authors:** Joshua Greig, Natalia A. Bulgakova

## Abstract

The regulation of E-cadherin at the plasma membrane by endocytosis is of vital importance for developmental and disease. p120-catenin, which binds to the E-cadherin C-terminus, can both promote and inhibit E-cadherin endocytosis. However, little is known about what determines the directionality of p120-catenin activity, and the molecules downstream. Here, we have discovered that p120-catenin fine-tunes the clathrin-mediated endocytosis of E-cadherin in *Drosophila* embryonic epidermal cells. It simultaneously activated two actin-remodelling pathways with opposing effects: RhoA, which stabilized E-cadherin at the membrane, and Arf1, which promoted internalization. Epistasis experiments revealed that RhoA additionally inhibited Arf1. E-cadherin was efficiently endocytosed only in the presence of intermediate p120-catenin amounts with too little and too much p120-catenin inhibiting E-cadherin endocytosis. Finally, we found that p120-catenin levels altered the tension of the plasma membrane. Altogether, this shows that p120-catenin is a central hub which co-ordinates cell adhesion, endocytosis, and actin dynamics with tissue tension.

## Introduction

Cell-cell adhesion is a fundamental requirement for the formation of tissues and organs. Such adhesions link cells to their neighbours and provide orientational information to a cell in a tissue [1]. In the epithelium, cell-cell adhesion is mediated by Adherens Junctions (AJs), with the principle component E-cadherin (E-cad), a transmembrane protein which binds in a homophilic fashion to E-cad molecules on adjacent cells [2,3]. Intracellularly E-cad interacts with the catenin protein family [4,5]. At its distal C-terminus E-cad binds β-catenin, with α-catenin binding β-catenin and actin, thus tethering extracellular adhesion with the cytoskeleton [4,6]. The third member, p120-catenin (p120ctn) binds to the JuxtaMembrane Domain (JMD) in the proximal C-terminus of E-cad [7–9]. These catenins are represented by a single gene for each in invertebrates including *Drosophila*, whereas both α-catenin and p120ctn families have several members in vertebrates. Thus, p120ctn family has 7 members in humans with different expression patterns and functional requirements, potentially reflecting an increase in tissue types and functional complexity [10–12].

The spatio-temporal expression and localization of E-cad is a prerequisite for the development of multi-cellular organisms: mutations of E-cad result in early embryonic lethality [13,14]. The loss of E-cad from the cell surface has been implicated in the progression of cancer cells towards metastatic spread [15]. Concurrently, the return of E-cad to the plasma membrane is required for cells to re-integrate into tissues to form secondary tumours [16,17]. Therefore, knowledge of the mechanisms that modulate E-cad levels at the cell surface is crucial for understanding regulation of E-cad in development and disease.

E-cad has multiple levels of regulation, with endocytic recycling providing a rapid response to changing tissue tension or dynamics [18,19]. The p120ctn family is as the key regulator of E-cad endocytosis in mammalian cells [20–26]. The most studies focused on the founding family member, p120ctn, however other members, δ-catenin and ARVCF seem to function in the same way and can substitute for p120ctn [27]. In mammalian cells, p120ctn is required to maintain E-cad at the plasma membrane: uncoupling p120ctn from E-cad or suppressing its expression results in complete internalization of E-cad [22,27,28], which has been reported in several cancers [29–31]. Thus, a “cap model” was proposed whereby the binding of p120ctn to E-cad concealed endocytosis triggering motifs [24,32]. This model of unidirectional p120ctn activity has recently been augmented in mammalian cells, when it was found that p120ctn promotes endocytosis of E-cad through interaction with Numb [25].

By contrast, in *Drosophila* and *C. elegans* p120ctn was thought less important playing only supporting role in adhesion as genetic ablation failed to replicate the effects observed in mammalian systems [33–35]. This is thought to be due to a greater similarity of invertebrate p120ctn to mammalian δ-catenin, ablation of which is similarly viable in mice [10,36]. However, δ-catenin expression is restricted to neural and neuroendocrine tissues [37], which is likely to explain the mildness of knockout phenotypes, whereas invertebrate p120ctn is broadly expressed in both epithelia and neurons [33], suggesting the potential functional similarity with mammalian p120ctn which shares the broad expression pattern [27]. Additionally, it has recently been reported that *Drosophila* p120ctn is required to stabilize E-cad in the pupal wing [38] and promotes the endocytosis and recycling of E-cad in the embryo and larval wing discs [39], indicating an evolutionary conservation of p120ctn function.

One key aspect of endocytic regulation of cell adhesion is the remodelling of the actin cytoskeleton by small GTPases, particularity of the cortical actin which lies parallel to the plasma membrane [40–42]. One of the best characterised of these GTPase regulators is RhoA, whose activity results in focal points of actin contraction. The interaction between Rho and E-cad has been well documented in mammalian systems, and is important for cancer cell progression and anoikis resistance of tumours [43]. In *Drosophila*, Rho promotes the regulated endocytosis of E-cad by Dia and AP2 [18]. Conversely, Rho activity antagonises endocytic events in the early embryo [44], indicating the pivotal role RhoA plays in E-cad endocytic dynamics. In mammalian cells, p120ctn can directly inhibit RhoA, indirectly inhibit it via p190RhoGAP, localize its spaciotemporal activity, or activate it [26,43,45–48], suggesting a context-dependent role of p120ctn in RhoA activity. p120ctn role in the RhoA pathway in *Drosophila* is unclear [49–51].

Another group of GTPases important in the endocytosis of E-cad are Arf (ADP-Ribosylating Family) proteins [52]. Arf GTPases are members of the Ras superfamily and recruit coat proteins to facilitate the intracellular trafficking of vesicles. The first member of the family, Arf1, is classically viewed as a Golgi resident and responsible for anterograde transport from the Golgi to the plasma membrane [53,54]. Recently, however, Arf1 was detected at the plasma membrane and participates in trafficking by co-operating with Arf6-dependent endocytosis [55,56]. Arf1 activity has been suggested to balance the initiation and impediment of endocytosis [44]. In *Drosophila*, Arf1 is required for the remodelling of the actin cytoskeleton and facilitating endocytosis in the early syncytial embryo [44,57,58]. Furthermore, Arf1 interacts with E-cad and another component of AJs, Par-3 [59,60].

Here, we demonstrate that p120ctn acts to both inhibit and promote endocytosis of E-cad within the same tissue. These activities are dependent on the amount of p120ctn at the plasma membrane and determine the levels and dynamic properties of E-cad. Further, we show that the interaction of p120ctn with Rho and Arf1 signalling pathways regulates clathrin-mediated endocytosis, with these two pathways acting in an opposing but coupled fashion to set the E-cad turnover rate. Additionally, we found that p120ctn determines tension at the tissue-level. Finally, we present a new model by which p120ctn fine-tunes adhesion through the regulation of actin dynamics and E-cad endocytosis, and thus modulates tissue tension.

## Results

### Both loss and overexpression of p120ctn stabilize E-cadherin at the plasma membrane via clathrin-mediated endocytosis

In previous work, in which the requirement of p120ctn to promote E-cad endocytosis in *Drosophila* was discovered [39], an E-cad expressed from a ubiquitous (*Ubi-p63E*) promoter was used (*Ubi∷E-cad*-GFP). This expression was in the presence of endogenous E-cad, which was untagged, thus potentially leading to E-cad overexpression which may be skewing the effects of p120ctn loss. We first sought to substantiate the previous findings using an E-cad tagged at its endogenous locus: *shotgun∷*E-cad-GFP (hereafter, E-cad-GFP). The cells of the epidermis of stage 15 *Drosophila* embryos exhibit a distinct rectangular morphology (Fig. 1A) with long cell borders which are orthogonal to the Anterior-Posterior axis of the embryo (hereafter referred to as the AP cell borders) and short borders which are orthogonal to the Dorsal-Ventral axis (DV cell borders). E-cad-GFP localizes in a narrow continuous band of fully formed mature AJs at these borders [61,62]. To measure protein amounts in this case and those described below, we quantified fluorescent intensities directly immitted by EGFP: an approached well-established in various models due to linear scaling of EGFP signal with the amount of protein present [63,64]. E-cad-GFP localized asymmetrically at this stage with a 1:2 (AP:DV) ratio between the two cell borders (Fig. 1A-C, Table S1) [65]. In the absence of both maternal and zygotic p120ctn, the amount of E-cad was reduced at both the AP and DV cell borders by approximately 15% (p=0.008 and p=0.035, respectively, Fig. 1A-C).

**Figure 1.**
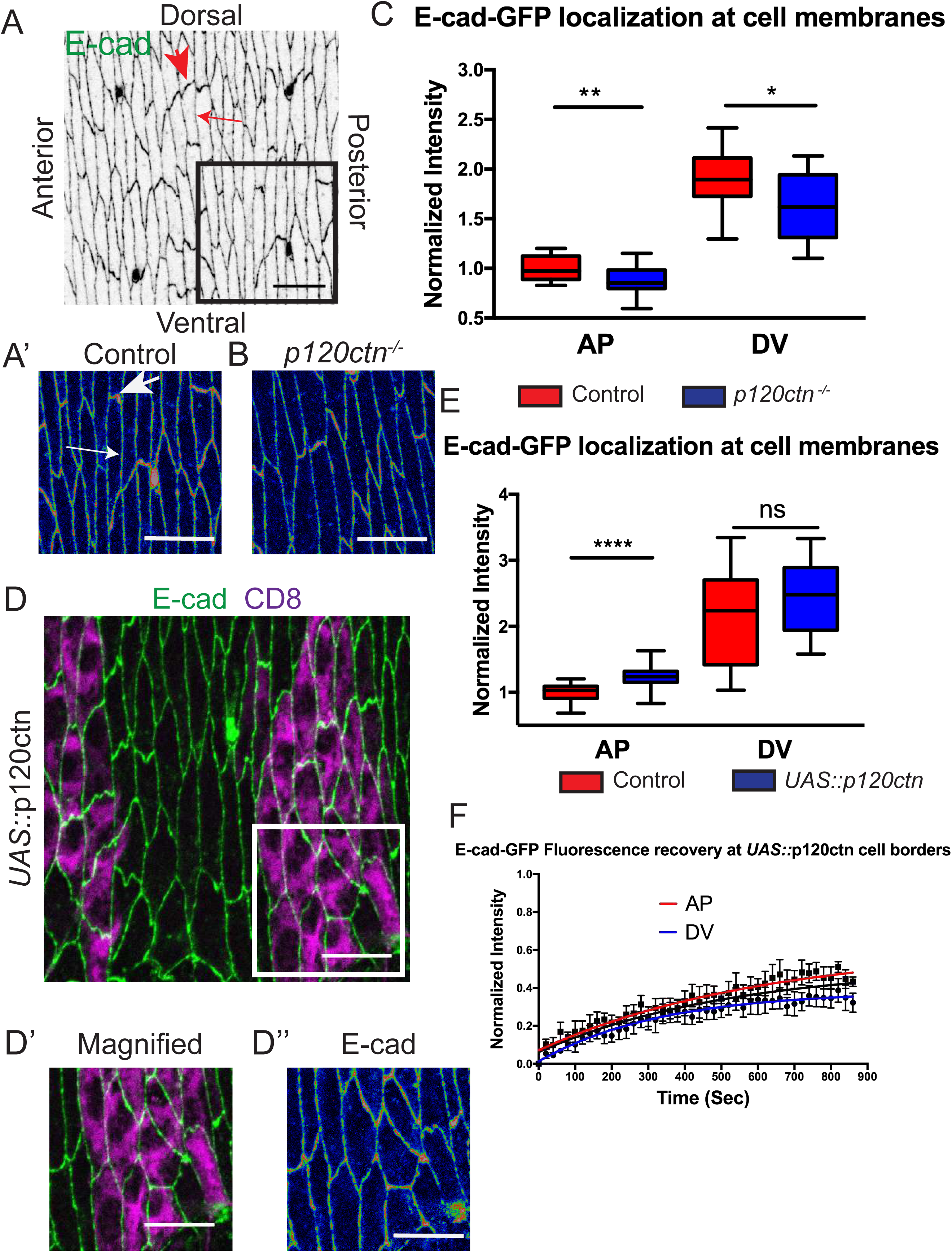
p120ctn levels determine the amount and dynamics of E-cad at the membrane. A-B. Apical view of the dorsolateral epidermis of stage 15 *Drosophila* wild type embryos (**A-A’**) and *p120ctn*^-/-^ mutant embryos (**B**) with cell outlines visualized by endogenously tagged E-cad-GFP using the rainbow spectrum for pixel intensity. Two distinct cell borders exist: the longer Anterior-Posterior (AP, large arrow) and the short Dorsal-Ventral (DV, small arrow). (**A’**) Magnified image of the epidermal cells indicated in the box of (A), scale bar is 10 µm. **C.** Quantification of the E-cad-GFP amount at the plasma membranes. **D.** Apical view of epidermis co-expressing *UAS∷*p120ctn with *UAS∷*CD8-Cherry using *en∷*GAL4 with E-cad-GFP (green in D-D’ and rainbow in D’’) and *UAS∷*CD8-Cherry (magenta in D-D’). *UAS∷*CD8-Cherry was used to mark cells expressing transgenes. (**D’-D’’**) magnified micrograph of the region indicated in (D), scale bar is 10 µm. **E.** Quantification of the E-cad-GFP amount at the plasma membrane in *UAS∷*p120ctn and internal control. **F.** Dynamics of E-cad-GFP measured by FRAP in the *UAS∷*p120ctn. Average recovery curves (mean ± s.e.m.) and the best-fit curves (solid lines) are shown. All best-fit and membrane intensity data are in Table S1. Bars in C and E represent difference between cell borders in genotypes measured by two-way ANOVA. *, P < 0.05; **, P < 0.01 ***, P < 0.001; ****, P < 0.0001. 10-20 embryos per genotype were used for intensity quantifications, and 8-10 embryos for FRAP.

Additionally, we measured the dynamics of E-cad-GFP using Fluorescence Recovery After Photobleaching (FRAP). In previous work, using the *Ubi∷*E-cad-GFP, recovery of the signal intensity at the membrane was 70% for DV and 50% for the AP plasma membranes in control. Recovery curves were best-fit by a bi-exponential model (Bulgakova *et al*., 2013), with the fast and slow recovery components attributed to diffusion and endocytic recycling, respectively [38,65,66]. The recovery was reduced to approximately 35% at both cell borders in *p120ctn* mutant embryos, resulting in an increase of immobile *Ubi∷*E-cad-GFP amounts at both junctions and therefore *Ubi∷*E-cad-GFP stabilization within AJs [39]. In the *p120ctn* mutant, a single exponential curve best described the data indicating that in the absence of p120ctn the slow recovery component observed in normal embryos was absent [39]. We replicated the above experiments using endogenously tagged E-cad-GFP, confirming that the effect of p120ctn loss was the same when E-cad was expressed at endogenous levels: the increase of the immobile fraction, and loss of the slow recovery component (Fig. S1, best-fit data in Table S1). Combining our results with published data, we conclude that p120ctn loss leads to the stabilization of E-cad at the membrane by affecting the slow endocytic component of recovery [39,65,67].

To further explore the function of p120ctn we examined the effect of its overexpression on E-cad. In mammalian cell culture, overexpression of p120ctn resulted in the elevation of VE-cad at plasma membranes by inhibiting endocytosis [68]. We created a p120ctn overexpression construct under the control of a *UAS* promoter (*UAS*∷p120ctn). We expressed it in the posterior half of each embryonic segment using the *engrailed*∷GAL4 (*en*∷GAL4) driver and marking the cells with a *UAS∷*CD8-Cherry (Fig. 1D). This resulted in an increase of E-cad-GFP at AP but not DV cell borders relative to the adjacent internal control cells (p=0.002 and p=0.27, respectively, Fig. 1E, Table S1). The result at the AP border was the reverse of that observed for the *p120ctn* mutant (Fig. 1C, Table S1). The absence of an increase at the DV might be due to E-cad being saturated at these borders in normal conditions. Measuring the dynamics of E-cad-GFP, using FRAP, we discovered that E-cad-GFP was less dynamic at both border types of the p120ctn overexpressing cells. Recovery was approximately 40% and 30% for AP and DV borders respectively (Fig. 1F), resulting in increased fraction of immobile E-cad-GFP at both borders (Table S1). Additionally, the slow recovery phase was lost, comparable to that observed in the case of p120ctn loss, indicating impairment of endocytic recycling (Table S1). The rate of fast recovery phase was the same in all cases. Therefore, both loss and overexpression of p120ctn impaired E-cad mobility and internalization.

Previously, the function of p120ctn in E-cad endocytosis was mostly studied by measuring the localization and internalization rate of E-cad (e.g. Davis et al., 2003; Pieters et al., 2016). We decided to directly examine the effect p120ctn has on components of the endocytic machinery and determine if p120ctn acts through clathrin-mediated endocytosis. We used Clathrin Light Chain (CLC) tagged with GFP (*UAS*∷CLC-GFP), to monitor clathrin localization and behaviour using light intensity released from the EGFP tag. CLC tagged with GFP incorporates functionally into clathrin-coated pits without interfering with basic clathrin functions in uptake, and is widely used to study dynamics of clathrin-mediated endocytosis [70–72], although there is some evidence that biochemically it is not identical to the untagged protein, it does reflect the physiological normal activity of clathrin [73]. CLC-GFP expressed using the *en*∷GAL4 driver was found in spots (puncta) localizing both on the plasma membrane in the plane of AJs and in the cytoplasm (Fig. 2A), a localization consistent with the known cellular function [74]. In the absence of p120ctn we observed a change in localization with more signal localizing with the E-cad at the membranes (Fig. 2B). Quantification of the puncta (see Materials and Methods) revealed that in the absence of p120ctn, CLC-GFP puncta were larger in area, fewer in number and more intense (p=0.002, p=0.0001, and p=0.0003, respectively, Fig. 2C-2E). As these are average measures of all puncta, we cannot distinguish if loss of p120ctn specifically increase CLC-GFP amounts and puncta size at the plasma membrane, or has an additional, likely indirect effect on puncta in cytoplasm. To explore the effect of p120ctn on clathrin dynamics we measured CLC-GFP recovery in the plane of the AJs using FRAP and found that CLC-GFP in the *p120ctn* mutant embryos was substantially less mobile, indicated by a smaller mobile fraction (p=0.0001, Fig. 2F).

**Figure 2.**
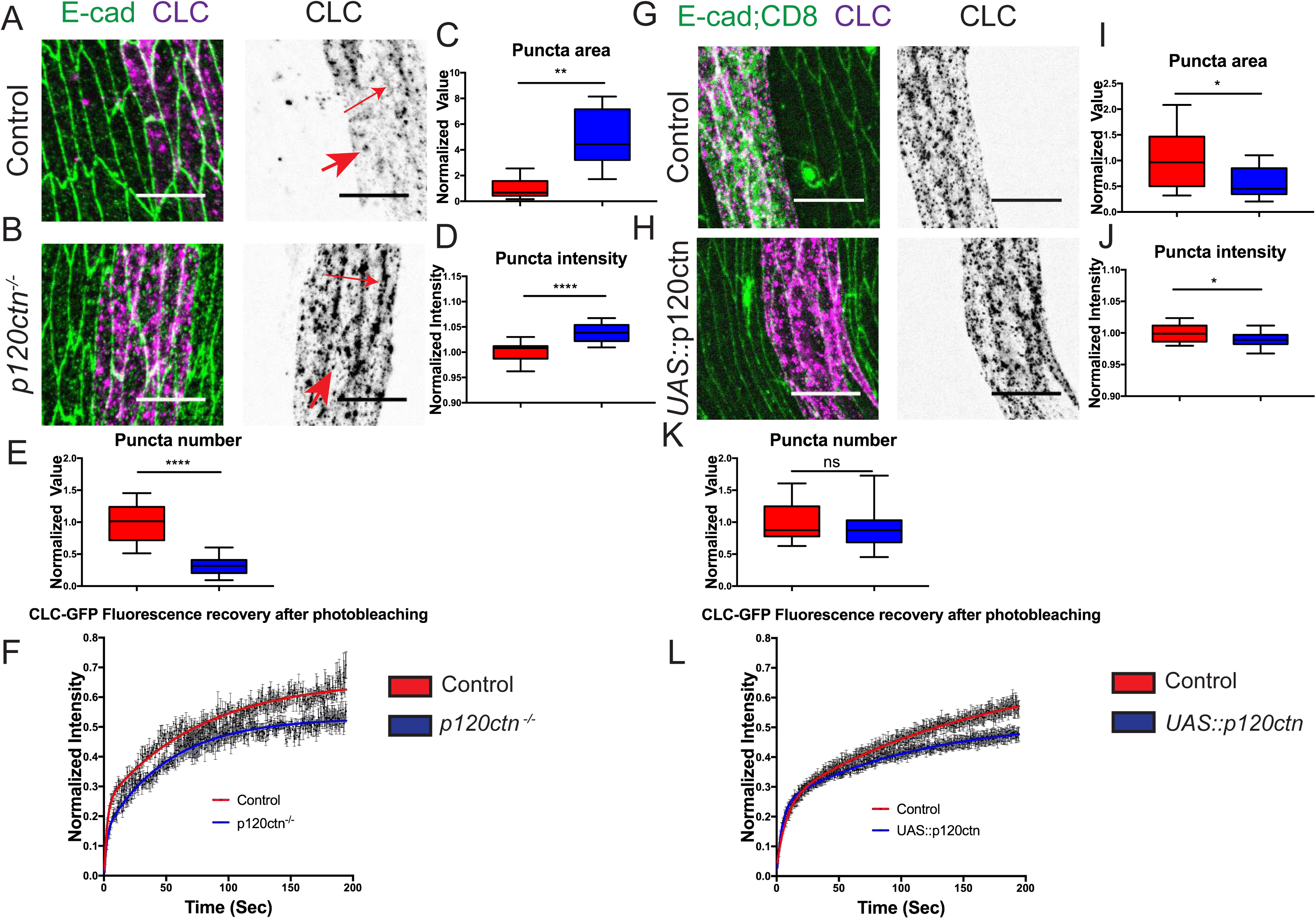
p120ctn regulation of E-cad is via the clathrin-mediated endocytic pathway. **A-B.** The localization of the clathrin light chain (*UAS∷*CLC-GFP) in p120ctn mutant (*p120ctn*^-/-^) embryos (**B**) and corresponding control (**A).** Distinct puncta (spots, magenta in left images and black in right images, indicated by arrows) are observed at the membrane and in the cytoplasm. Cell outlines are visualized by anti-E-cad antibody staining (green in left images). **C-E.** Quantification of the clathrin puncta in p120ctn mutant (*p120ctn*^-/-^) embryos by measuring the area (size, **C**), intensity (**D**), and the number (**E**) in control and p120ctn mutant. **F.** FRAP of *UAS*∷CLC-GFP in p120ctn mutant embryos with average recovery curves (mean ± s.e.m.) and the best-fit curves (solid lines). **G-K.** The localization of the clathrin light chain (*UAS∷*CLC-GFP) in embryos overexpressing p120ctn (*UAS∷*p120ctn) (**G**) and corresponding control (**H).** Distinct puncta (spots, magenta in left images and black in right images) are observed at the membrane and in the cytoplasm. Cell outlines are visualized by anti-E-cad antibody staining (green in left images). Note that in control for p120ctn overexpression, there is additionally signal from *UAS*∷CD8-mCherry in green. Quantification of the clathrin puncta in p120ctn overexpression (*UAS∷*p120ctn) embryos by measuring the area (size, **I**), intensity (**J**), and the number (**K**) in control and p120ctn overexpression. **L.** FRAP of *UAS*∷CLC-GFP in *UAS∷*p120ctn embryos with average recovery curves (mean ± s.e.m.) and the best-fit curves (solid lines). All best-fit and membrane intensity data are in Table S1. Statistical analysis is a two-tailed students t-test with Welch’s correction. *, P < 0.05; **, P < 0.01 ***, P < 0.001; ****, P < 0.0001. 10-20 embryos per genotype were used for puncta quantifications, and 8-10 embryos – for FRAP.

As E-cad had a smaller mobile fraction in p120ctn overexpression (Fig. 1F) we sought to determine whether this was due to an effect on the clathrin-mediated endocytic pathway. To explore this, we overexpressed p120ctn in conjunction with CLC-GFP. To balance the GAL4-UAS copy number, we co-expressed CD8-Cherry with CLC-GFP in the control (Fig 2G. CLC-GFP puncta in the p120ctn overexpressing cells were smaller in area and less intense (p=0.01 and p=0.03, respectively, Fig. 2H-2J), and overall puncta number was unchanged (Fig. 2M), indicating a probable defect in the formation of mature vesicles which abscise from the membrane. Finally, measuring the dynamics of CLC-GFP in these cells revealed that p120ctn overexpression reduced the mobile fraction of CLC-GFP (p=0.0001, Fig. 2N). Therefore, as both the removal and overexpression of p120ctn resulted in reduction of clathrin mobility, we reasoned that p120ctn levels act in a bell-curve fashion to regulate clathrin-mediated endocytosis of E-cad.

### p120ctn acts via the Rho signalling pathway to stabilize E-cad at the adherens junctions

A primary candidate to link p120ctn to clathrin-mediated endocytosis is the GTPase RhoA. To ascertain whether p120ctn acts on clathrin-mediated endocytosis through regulating Rho signalling we turned to the downstream effectors of RhoA: the enzyme Rho-Kinase (Rok), which specifically binds RhoA in its activated form, resulting in Rok recruitment to membranes [75]; and the actin cross-linker non-muscle Myosin II, a Rho-kinase target which requires Rho-kinase for recruitment to AJs [76–78].

We used a tagged kinase-dead variant of Rok (Venus-Rok^K116A^, hereafter referred to as Rok-Venus) driven by the *spaghetti squash* promoter as an established readout for the normal localization of Rok without overactivation of the signalling pathway [79]. In the epidermis we observed a distinct asymmetry of Rok-Venus localisation between the AP and DV cell borders in the order 2:1 (AP:DV, Fig.3A-D) consistent with previous reports [65,79]. In the *p120ctn* mutant embryos we detected a loss of this asymmetry, specifically due to a reduction of the Rok-Venus levels at the AP cell borders (p=0.013, Fig. 3A-C). To confirm the interaction between p120ctn and Rok, we examined the effect of p120ctn overexpression. We found an increase in the amount of Rok-Venus at the AP cell borders (p=0.0043, Fig. 3A-B,D). Therefore, the levels of Rok-Venus at the plasma membrane positively change with those of p120ctn.

**Figure 3.**
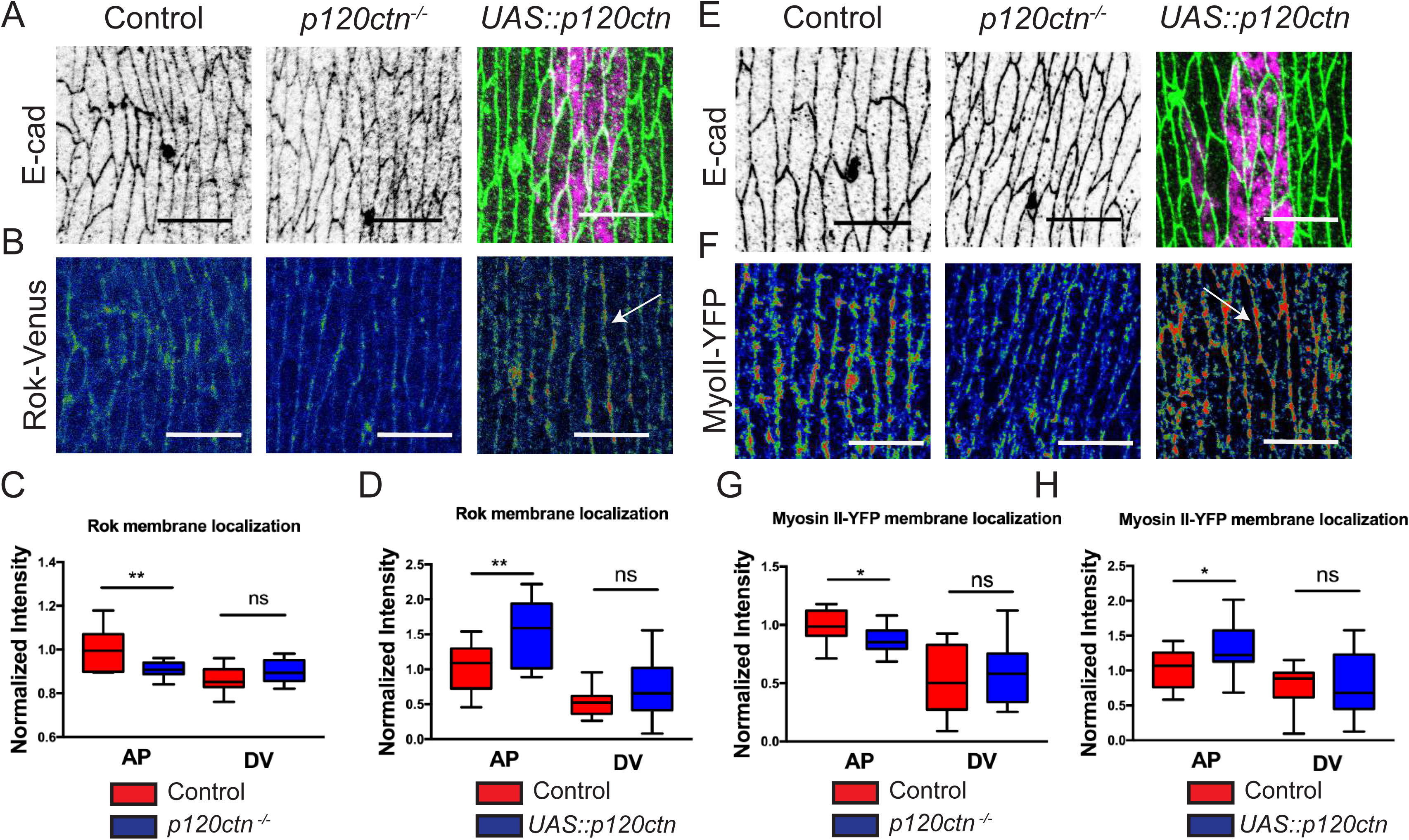
p120ctn activates RhoA signalling at the plasma membrane. **A-B.** Apical view of the epidermis of control, p120ctn mutant (*p120ctn*^-/-^) and p120ctn overexpression (*UAS∷*p120ctn) embryos visualized with anti-E-cad antibody (**A**, grey in the left and middle images, green in the right image), *UAS*∷CD8-mCherry (**A**, magenta in the right image, marks cells expressing *UAS∷*p120ctn) and Rho-Kinase (Rok) tagged with Venus (**B,** rainbow pixel intensity spectrum, membrane indicated by arrow). **C-D.** Quantification of Rok-Venus in the p120ctn mutant (**C**) and p120ctn overexpressing embryos (**D**). **E-F.** The epidermis of control, p120ctn mutant (*p120ctn*^-/-^) and p120ctn overexpression (*UAS∷*p120ctn) embryos visualized with anti-E-cad antibody (**E**, grey in the left and middle images, green in the right image), *UAS*∷CD8-mCherry (**E**, magenta in the right image, marks cells expressing *UAS∷*p120ctn), and Myosin II (MyoII) tagged with YFP (**F,** rainbow pixel intensity spectrum, membrane indicated by arrow). **G-H.** Quantification of MyoII-YFP in the p120ctn mutant (**G**) and p120ctn overexpressing embryos (**H**). All membrane intensity data are in Table S1. Scale bars – 10 µm. Upper bars in **C, D, G** and **H** represent difference between genotypes and the lower - between cell borders measured by two-way ANOVA. *, P < 0.05; **, P < 0.01 ***, P < 0.001; ****, P < 0.0001. 12-20 embryos per genotype were quantified.

To directly demonstrate an effect of p120ctn on the cytoskeleton we used an endogenously tagged variant of the non-muscle Myosin II (MyoII-YFP), which reproduces the localization of untagged Myosin II [78]. The localization of the MyoII-YFP protein corresponded to that of Rok-Venus (Fig. 3E-F). The loss of p120ctn resulted in the same effect on MyoII-YFP as that observed on Rok-Venus: loss of asymmetry due to a reduction at the AP cell borders (p=0.0011, Fig. 3E-G). Similarly, p120ctn overexpression resulted in an increase of MyoII-YFP at the AP plasma membrane by comparison to the internal control (p=0.025, Fig. 3E-F,H). Together, these results support a mechanism whereby p120ctn activates Rho signalling in a dose-dependent manner.

Having established an interaction between RhoA and p120ctn, we next examined the impact on E-cad endocytosis using constitutively active (Rho^CA^) and dominant negative (Rho^DN^) constructs to modulate Rho signalling. Dampening of Rho activity using suppression of the activator RhoGEF2 reduced E-cad levels at plasma membranes [65]. As this was a mild suppression by RNAi we decided to measure the effect of expressing Rho^DN^. Expressed using the strong driver *en∷GAL4*, it resulted in a complete loss of E-cad at the membrane by stage 15 of embryogenesis (data not shown). Therefore, we acutely induced the expression of the Rho^DN^ using temperature sensitive GAL80^ts^ expressed from the *tubulin84B* promoter [80]. We measured the amount of E-cad-GFP at the plasma membranes four hours post-induction of the Rho^DN^ expression, with time adjusted to obtain stage 15 of embryogenesis. In this case, cells presented the residual E-cad-GFP at the plasma membranes but, strikingly, it was reduced to discrete puncta along the membrane (arrowhead, Fig. 4A). The levels of E-cad-GFP at the plasma membranes were reduced at the AP cell borders (p=0.0039, Fig. 4B).

**Figure 4.**
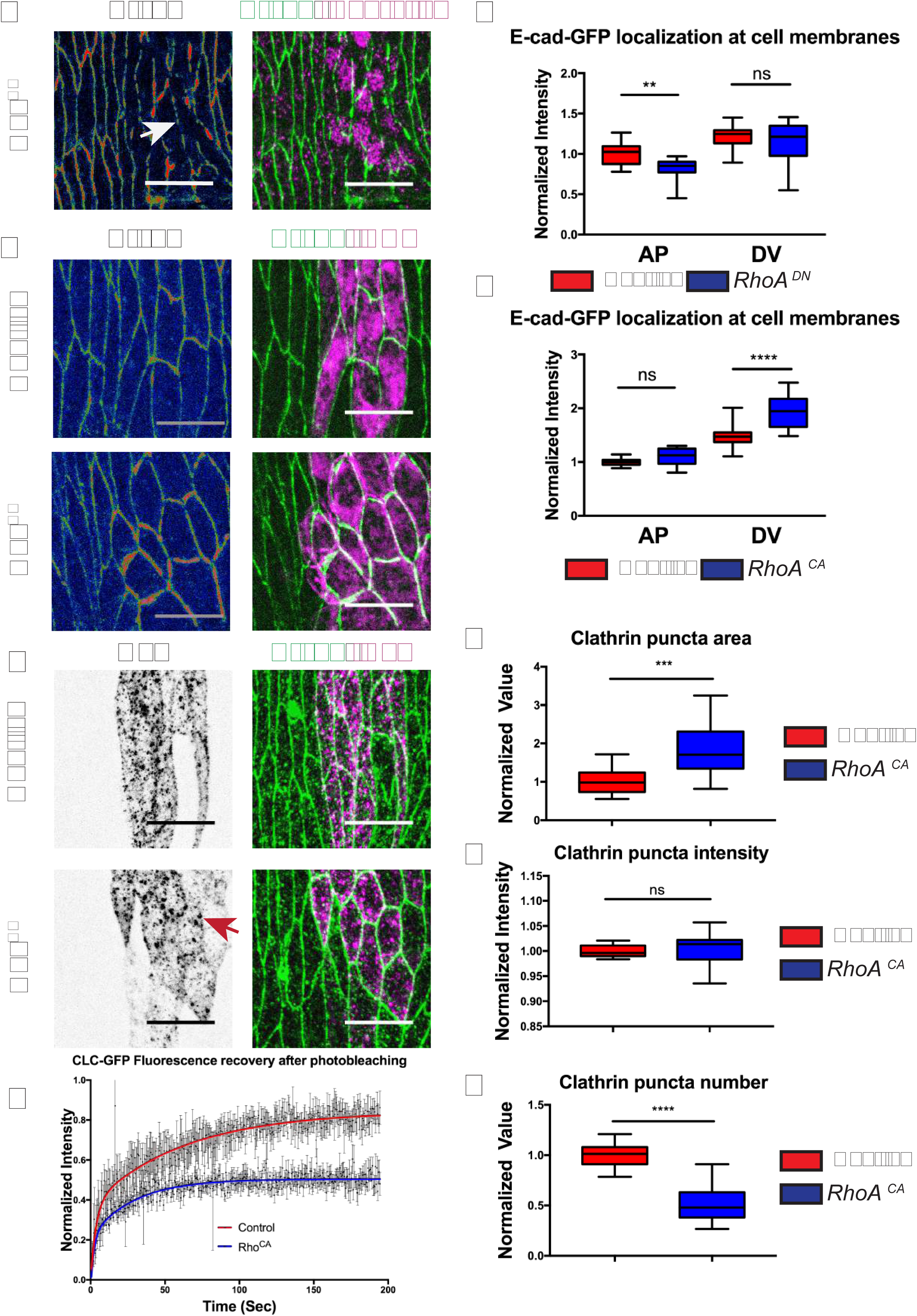
RhoA activity promotes the stabilization, and opposes the internalization, of E-cad at the membrane. **A-D.** Localization of E-cad in the epidermis following downregulation (the induction of Rho^DN^ expression for 4 hours, **A-B,** membrane indicated by arrow) or upregulation (expression of Rho^CA^, **C-D**, bottom images in **C** with top images being the corresponding control expressing *UAS*∷CD8-Cherry alone) of Rho signalling. Cell borders are marked by E-cad-GFP (rainbow spectrum in left images, green in right images). Cells expressing transgenes are marked with an antibody for Engrailed directly (**A**) or by *UAS∷*CD8-Cherry (**C**). Quantification of the E-cad-GFP at the membrane of the Rho^DN^ (**B**) or Rho^CA^ (**D**) expressing cells in comparison to internal control cells (**B**) or external controls (**D**). The total amount of E-cad-GFP per cell upon upregulation of Rho signalling by Rho^CA^ expression is in Fig. EV3. **E.** Localization of clathrin (*UAS*∷CLC-GFP, grey in left images, magenta in right images) in cells co-expressing Rho^CA^ (bottom) and corresponding control co-expressing *UAS*∷CD8-Cherry (top). Cell borders are visualized by anti-E-cad antibody staining (green in right images, arrow indicates membrane localized clathrin puncta). **F-H.** Quantification of the clathrin puncta area (F), intensity (G), and number (H) in the Rho^CA^ expressing cells. **I.** FRAP of *UAS*∷CLC-GFP in Rho^CA^ expressing cells. Average recovery curves (mean ± s.e.m.) and the best-fit curves (solid lines) are shown. All best-fit and membrane intensity data are in Table S1. Scale bars – 10 µm. *, P < 0.05; **, P < 0.01 ***, P < 0.001; ****, P < 0.0001. 10-20 embryos per genotype were used for puncta quantifications, and 8-10 embryos – for FRAP.

To complement these data, we examined the effect of overactivation of Rho signalling using the Rho^CA^ construct. This construct resulted in stark changes in cell morphology (Fig. S2) and an increase of E-cad-GFP membrane levels at DV cell borders (p<0.0001, Fig. 4C-D). Due to cells rounding up, thus complicating the division into true AP and DV cell borders, we also measured the mean intensity of E-cad-GFP at the membrane of these cells as a total of the entire membrane rather than dividing into AP and DV (Fig. S2). As expected this analysis supported the previous result: E-cad-GFP intensity was significantly increased in the Rho^CA^ cells (p<0.0001, Fig. S2). Overall, these results showed that RhoA activity positively correlated with E-cad localization at the plasma membrane.

To investigate if the observed effects of manipulating Rho activity were the result of an endocytic mechanism, we measured the effects of these Rho constructs on clathrin using CLC-GFP. Co-expression of CLC-GFP with Rho^CA^ resulted in a localization change of the clathrin puncta in comparison to co-expression with CD8-Cherry in the control (Fig. 4E). A larger proportion of protein was localizing with E-cad at the membrane, and the puncta were of greater area and fewer in number (p=0.0009 and p<0.0001, respectively, Fig. 4F, 4H). Finally, we measured the dynamics of clathrin in these cells. The recovery of CLC-GFP was lower (p<0.0001, Fig. 4I), suggesting that Rho activity impairs the budding of clathrin coated vesicles, leading to retention at the plasma membrane. In conclusion, our data is consistent with p120ctn-dependent activation of Rho signalling resulting in the inhibition of clathrin mediated endocytosis, enabling p120ctn to stabilize E-cad at the cell surface.

### p120ctn promotes the internalization of E-cadherin through Arf1 signalling

Due to the reported interactions of Arf1 with E-cad and Par-3 [59,60] we decided to examine if Arf1 acts downstream of p120ctn. We used a transgenic GFP tagged variant of Arf1 (*UAS*∷Arf1-GFP) driven by *en∷*GAL4. Although Arf1 tagged with GFP has reduced affinity for ArfGAPs and ArfGEFs, as well as nucleotide exchange rate [81], this might be beneficial for the experiments described below, as it allows the study of Arf1 without hyperactivating the pathway. Expressing this transgene in the epidermis we observed large and distinct puncta in the cytoplasm including in proximity to the plasma membrane, which corresponding to the Golgi-resident Arf1 population (Fig. 5A, Fig. S3). To lesser extent, Arf1 localized at the plasma membranes in the plane of AJs, similarly to what was reported before [59] (Fig. 5A, highlighted by arrows). As we were interested in the localization of Arf1 at the plasma membrane not the Golgi we excluded the puncta signal from our quantification (see Materials and Methods). Removing p120ctn resulted in a decrease in the amount of Arf1-GFP at the membrane, affecting both the AP and DV cell borders (p<0.0001 and p<0.0001, respectively, Fig. 5A-C). In a complementary experiment, we overexpressed p120ctn: in these cells the amount of Arf1-GFP at the membrane was reduced at the AP border plasma membranes (p=0.02, Fig. 5D-F). Therefore, both the loss and overexpression of p120ctn reduced the amount of Arf1 at the membrane. This was highly reminiscent of the result we observed for clathrin. Moreover, as Arf1 is a known recruiter of clathrin at the Golgi [82], we inferred that plasma membrane-resident Arf1 is downstream of p120ctn, which may link it to the clathrin-mediated endocytic machinery.

**Figure 5.**
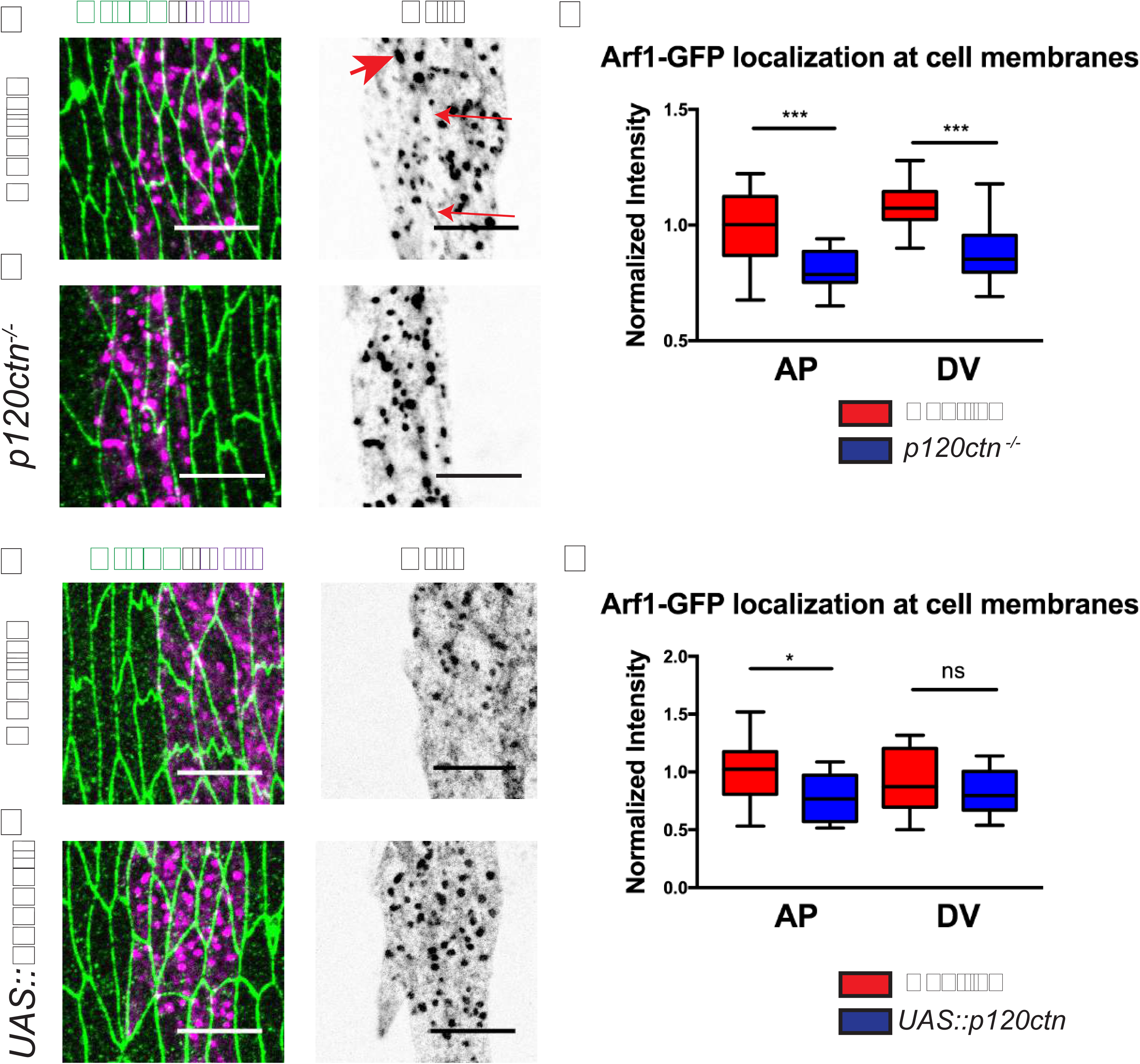
Arf1 is a downstream interactor of p120ctn. **A-B.** Apical views of the epidermis expressing *UAS∷*Arf1-GFP (Arf1, magenta in left images and grey in right images) with *en*∷Gal4 in p120ctn mutant (*p120ctn*^-/-)^, **B**) and corresponding control (**A**). Cell outlines are visualized with anti-E-cad antibody staining (green in left images). The large *UAS*∷Arf1-GFP puncta in the cytoplasm (large arrow in A) marks the Golgi (see Fig. EV3). The small arrow in A indicates the membrane resident population of Arf. **C.** Quantification of the amount of *UAS∷*Arf1-GFP at the plasma membrane between a control and p120ctn mutant embryos. **D-E.** Apical views of the epidermis expressing *UAS∷*Arf1-GFP (Arf1, magenta in left images and grey in right images) in embryos overexpressing p120ctn (*UAS*∷p120ctn, **E**), and corresponding controls (**D**). Cell outlines are visualized with anti-E-cad antibody staining (green in left images). **F.** Quantification of the amount of *UAS∷*Arf1-GFP at the plasma membrane between a control and p120ctn overexpressing embryos. All membrane intensity data are in Table S1. Scale bars – 10 µm. Upper bars in **C** and **F** represent difference between genotypes and the lower - between cell borders measured by two-way ANOVA. *, P < 0.05; **, P < 0.01 ***, P < 0.001; ****, P < 0.0001. 10-20 embryos per genotype were quantified.

Having determined an interaction between p120ctn and Arf1, we next asked whether Arf1 activity had a more directive role in p120ctn mediated endocytosis of E-cad. Expression of a dominant negative variant of Arf1 (Arf1^DN^, Wang et al., 2017) using the *en*∷GAL4 driver increased the amount of E-cad-GFP at the plasma membrane at the AP borders with no change detected at the DV (p=0.02 and p=0.5, respectively Fig. 6A-B). This was accompanied by an abnormal cell morphology (Fig. 6A and Fig. S4). No surviving larvae were observed, consistent with previous reports using Arf1^DN^ in *Drosophila* [84]. We attributed this lethality to perturbation of post-Golgi protein transport by prolonged exposure to Arf1^DN^, leading to cell death [81,85].

**Figure 6.**
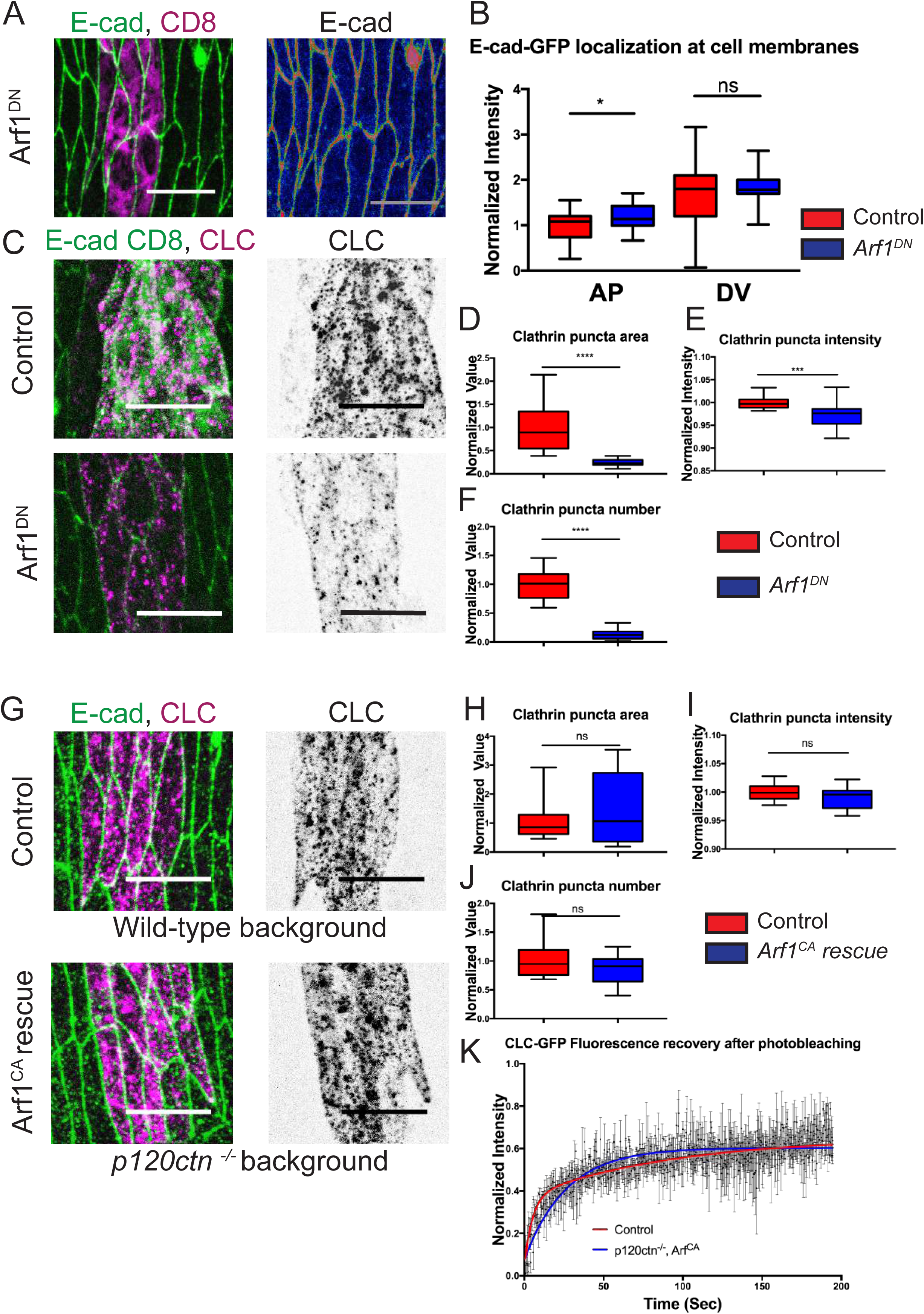
Arf1 activity promotes the clathrin-mediated internalization of E-cad, and Arf1 overactivation rescues the defects in clathrin localization and dynamics in p120ctn mutant background. **A.** Localization of E-cad (E-cad-GFP, green in left image, and rainbow in right image) in cells expressing a dominant negative variant of Arf1 (Arf1^DN^) with *en*∷GAL4. Cell borders are visualized with E-cad-GFP. *UAS*∷CD8-Cherry was used to mark cells expressing Arf1^DN^ (magenta, left image). **B.** Quantification of the amount of E-cad-GFP at the plasma membranes of the Arf1^DN^ cells compared to adjacent internal control cells. **C.** Localization of *UAS*∷CLC-GFP (magenta in left images, and grey in right images) in cells expressing a dominant negative variant of Arf1 (Arf1^DN^) with *en*∷GAL4. Note that in control there is additionally signal from *UAS*∷CD8-mCherry in green. Cell borders are visualized with anti-E-cad antibody staining (C, left image). *UAS*∷CD8-Cherry was used to balance the dose of GAL4-UAS (green, top images). **D**-**F.** Quantification of the *UAS*∷CLC-GFP puncta area (**D**), intensity (**E**), and number (**F**) in the plane of AJs in the Arf1^DN^ cells. **G.** Localization of *UAS*∷CLC-GFP (magenta in left images, grey in right images) in *wt* control, expressing *UAS∷*CLC-GFP alone (top images), and in embryos which are expressing a constitutively active Arf1 (Arf1^CA^) in a p120ctn mutant (*p120ctn*^-/-^) genetic background (bottom images). Cell borders were visualized with anti-E-cad antibody staining (green in right images). **H-J.** Quantification of the *UAS∷*CLC-GFP puncta area (**H**), intensity (**I**), and number (**J**) in the plane of AJs in the Arf1^CA^; *p120ctn*^-/-^ embryos. **K.** FRAP of *UAS∷*CLC-GFP in the plane of AJs in the Arf1^CA^; *p120ctn*^-/-^ embryos with average recovery curves (mean ± s.e.m.) and the best-fit curves (solid lines). All best-fit and membrane intensity data are in Table S1. Scale bars – 10 µm. *, P < 0.05; **, P < 0.01 ***, P < 0.001; ****, P < 0.0001. 10-20 embryos per genotype were used for puncta quantifications, and 8-10 embryos – for FRAP.

To ascertain if Arf1 was acting on E-cad membrane localization via clathrin we measured the effect of expressing Arf1^DN^ on the localization and amount of CLC-GFP. Co-expression of CLC-GFP with Arf1^DN^ resulted in a substantial reduction in CLC-GFP puncta area, intensity, and number (p<0.0001, p=0.0005, and p<0.0001, respectively, Fig. 6C-F). Therefore, we conclude that Arf1 is required for the normal recruitment and function of clathrin at the AJs.

Finally, to confirm that Arf1 is functionally downstream of p120ctn we designed a rescue experiment using a constitutively active Arf1 (Arf1^CA^). We expressed this in a *p120ctn* mutant background and measured the effect on CLC-GFP (see Fig. 2). We compared this to control co-expressing CLC-GFP with CD8-Cherry in an otherwise wild-type genetic background. Control clathrin puncta localization was as previously observed (Fig. 6G). In the *p120ctn* mutant expressing Arf1^CA^ the clathrin puncta were no different in area, intensity or number from control (p=0.21, p=0.19, and p=0.33, respectively, Fig, 6H-J). Measuring the dynamics of clathrin revealed that recovery was no longer significantly different from the wild-type control (Fig. 6K). From both fixed and dynamic measures of clathrin we concluded that the expression of Arf1^CA^ rescues the clathrin defect observed in the *p120ctn* mutant (see Fig. 2). Therefore, Arf1 functionality is consistent with it being a downstream interactor of p120ctn, which links the p120ctn–E-cad complex to the clathrin-mediated endocytic machinery. Overall, this allows p120ctn to promote the internalization of E-cad in addition to the previously described anti-endocytic function with RhoA signalling.

### The Rho signalling pathway is upstream of Arf1

Having established that Arf1 and RhoA signalling are regulated by p120ctn, resulting in the internalization or retention of E-cad respectively, we sought to determine if these two pathways act upon one another or exist independently. As Arf1 was reported to regulate RhoA [86], we explored the possibility of an interaction between Arf1 and Rho signalling. To address this, we perturbed signalling in one pathway and measured the effect on the other.

First, we measured the membrane localization of MyoII-YFP, as a readout of RhoA signalling activity, upon upregulation of Arf1 signalling using Arf1^CA^. MyoII-YFP localization was indistinguishable between cells expressing Arf1^CA^ and control cells (Fig. 7A-B), demonstrating that Rho signalling in embryonic epidermis is independent of Arf1 function. In a complementary experiment, we impaired RhoA signalling using an RNAi against the upstream activator RhoGEF2, which results in a milder E-cad reduction at AJs than the severe loss observed with Rho^DN^ expression (Fig. 4 and Bulgakova et al., 2013). We measured the membrane levels of Arf1-GFP in cells additionally expressing either RhoGEF2-RNAi or CD8-Cherry (Fig. 7C, D). Downregulation of RhoGEF2 resulted in a significant increase in the amount of Arf1-GFP at both AP and DV borders (p=0.045 and p=0.022, respectively, Fig. 7E), demonstrating that RhoA signalling activity limits Arf1 localization to the plasma membrane. This was reminiscent of the phenotype observed upon p120ctn overexpression, in which Arf1-GFP is reduced at the plasma membrane (see Fig. 5). Therefore, as the overexpression of p120ctn is accompanied by upregulation of Rho signalling (see Fig. 3), we suggest that the reduction of Arf1 previously observed (Fig. 5) is caused by the regulation of Arf1 recruitment by Rho signalling. However, the reduction of Arf1 at the plasma membrane in the absence of p120ctn (see Fig. 5) is independent of RhoA and is caused by regulation of Arf1 recruitment and/or activation more directly by p120ctn.

**Figure 7.**
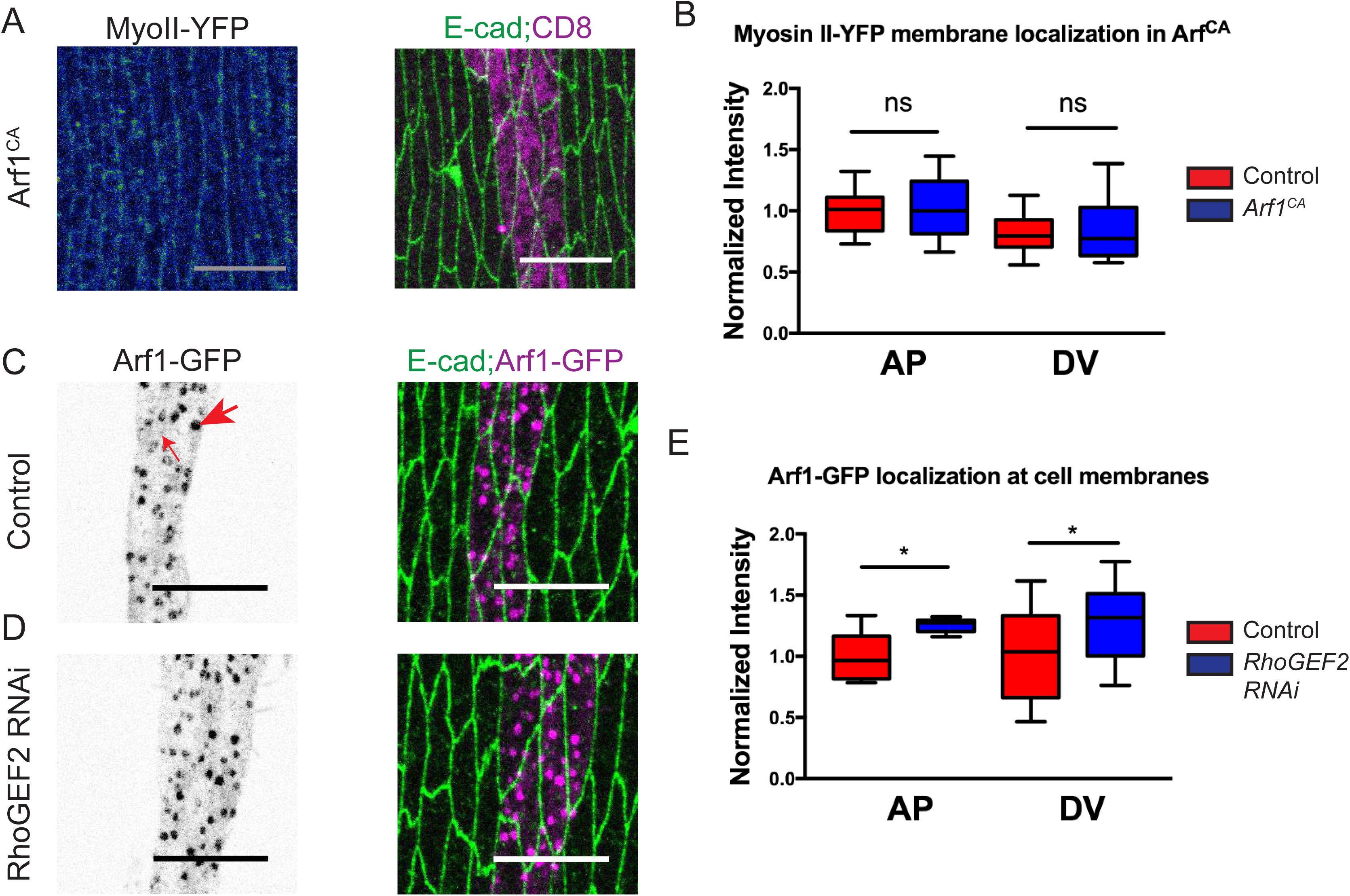
RhoA signalling inhibits Arf1 and is independent of it. **A.** Localization of MyosinII-YFP (rainbow in left image) in the epidermis of cells expressing a constitutively active Arf1 (Arf1^CA^) with *en*∷GAL4 cells marked by *UAS∷*CD8-Cherry (magenta in right image). Cell borders are visualized with anti-E-cad antibody staining (green in right image). **B.** Quantification of the MyoII-YFP at the plasma membranes of the Arf1^CA^ cells and the adjacent internal control plasma membranes. **C**-**D.** The localization of *UAS∷*Arf1-GFP (grey in left images, magenta in right images) in control (**C)** and cells expressing RhoGEF2 RNAi (**D**). Large arrow indicates the Golgi resident Arf1 and the small arrow indicate the membrane resident population in **(C)**. Cell borders were visualized with anti-E-cad antibody staining (green in right images). **E.** Quantification of the amount of *UAS∷*Arf1-GFP at the plasma membranes of the RhoGEF2RNAi expressing cells. All membrane intensity data are in Table S1. Scale bars – 10 µm. *, P < 0.05; **, P < 0.01 ***, P < 0.001; ****, P < 0.0001. 10-20 embryos per genotype were quantified.

### p120ctn levels modulate plasma membrane tension

Having identified two actin-remodelling pathways downstream of p120ctn we sought to understand their functional outcome at the tissue-level. Actin assembly on clathrin-coated pits, which requires Arf activity [87], is suggested to counteract cortical tension to enable membrane deformation [88]. In light of the MyoII results (see Fig. 3E-F), we decided to test if p120ctn modulated membrane tension. We analysed this using microablation of membranes and measured the initial recoil as a readout of tissue tension [89]. We examined AP membranes as these presented an enrichment of MyoII (see Fig. 3E-F). The initial recoil is proportionally influenced by the underlying tension in the system rather than other variables [90], thus we decided to focus our attention on this measure. The initial recoil was measured as the distance between the two connected cell vertices of the ablated membrane immediately post-ablation, expressed as a change in the length proportional to pre-ablation length (Fig. 8A). In control embryos expressing E-cad-GFP alone, the recoil distance post-ablation showed an increase of 5% over pre-ablation distance (Fig. 8A-B). The overexpression of p120ctn resulted in a higher mean initial recoil of 10% over the pre-ablation distance (p<0.0001, Fig 8A-B). Conversely, in *p120ctn* mutant cells the recoil was decreased (p=0.022) to a mean value of 2% increase over pre-ablation distance (Fig. 8A-B). Therefore, membrane tension positively correlated with p120ctn.

**Figure 8.**
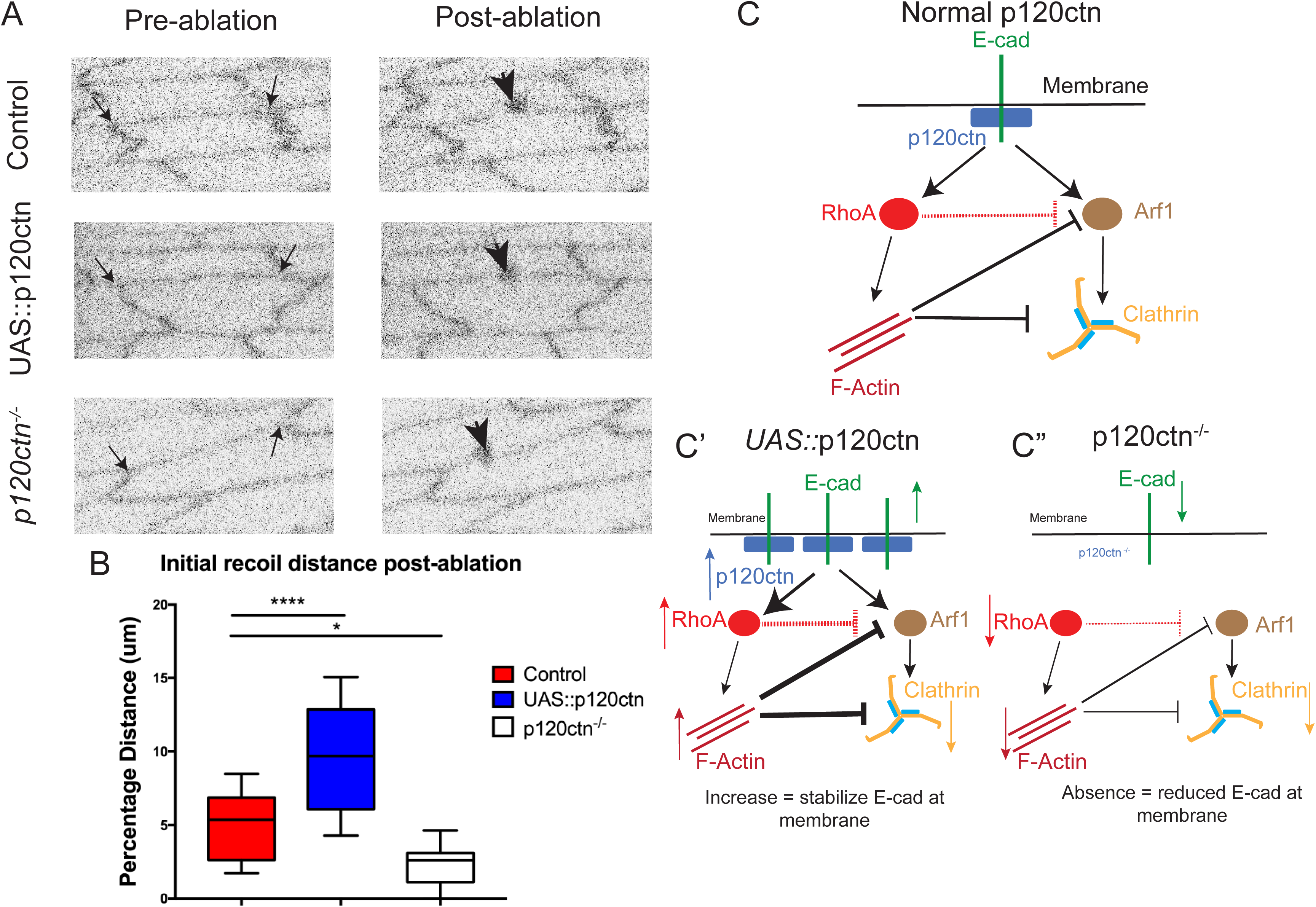
p120ctn increases the tension at the plasma membrane, and the model of p120ctn action. **A.** Examples of laser ablation in epidermal cells in control (top images), p120ctn mutant (*p120ctn*^-/-^, middle images), and p120ctn overexpressing (*UAS*∷p120ctn, bottom images). Cells shown immediately before (left images) and after ablation (right images). Cell borders are visualized with E-cad-GFP (grey). The small arrows indicate the connected vertices of the ablated membrane used to measure distance during the course of the experiment. The large arrows in the post-ablation images represent the position at which the microablation was performed, for which a small spot of discolouration can be observed. **B.** Quantification of the initial recoil of the ablated membranes, expressed as a percentage increase in distance relative to the length of the membrane prior to ablation. **C.** Model of the interaction between p120ctn, RhoA, Arf1, clathrin, and actin. In normal conditions p120ctn acts on both Arf1 and RhoA signalling pathways: Rho promotes the stabilization of E-cad at the plasma membrane and Arf1 promotes E-cad internalization by recruiting clathrin. RhoA additionally inhibits Arf1, possibly directly (dashed red lines) or indirectly by increasing the density of the cortical actin. (**C’**) Overexpression of p120ctn leads to overactivation of the Rho signalling pathway, resulting in an increase in cortical actin and indirect suppression of Arf1 signalling. Overall this results in an increase in E-cad amount and stability at the membrane. (**C”**) Conversely in the absence of p120ctn both the Rho and Arf1 pathways are not stimulated. Therefore, both cortical actin tension and clathrin mediated endocytosis are reduced. The lower levels of E-cad at the membrane result from the reduction of recycling and actin mediated stabilization by p120ctn. Scale bar – 10µm. *, P < 0.05; **, P < 0.01 ***, P < 0.001; ****, P < 0.0001. 14-20 embryos per genotype.

## Discussion

In this study we have focused on the mediators and mechanism of E-cad turnover downstream of p120ctn. We present several advances to the understanding of p120ctn activity and its role in cell-cell adhesion. First, we showed that p120ctn acts bi-directionally as a set point for E-cad at the plasma membrane via a clathrin-mediated mechanism. Second, we demonstrated that p120ctn regulates RhoA signalling, which regulates the internalization of E-cad. This places *Drosophila* p120ctn in the same context as its mammalian orthologue, which interacts with Rho [91]. Third, we discovered an Arf1 dependent mechanism by which p120ctn promotes the internalization of E-cad. The described changes in protein amounts following manipulation of p120ctn were in the range of 10%-40%, which may be considered subtle and would be in line with the notion that p120ctn has supportive rather than essential role in non-chordate adhesion [33,35]. We speculate that the mildness of changes is due to existence of redundancies and feedback interactions to ensure robustness of cadherin adhesion, essential in a multicellular organism [92]. Altogether, we propose a new model of p120ctn action, in which the activity of p120ctn as a function of the differential interaction with RhoA and Arf1 (Fig. 8C). In this model, p120ctn regulates membrane tension through RhoA, reinforcing cortical actin, which inhibits E-cad endocytosis. Simultaneously, p120ctn regulates the formation of clathrin-coated vesicles through Arf1, promoting E-cad endocytosis, until a threshold is passed and further increase of p120ctn results in Arf1 inhibition, due to the concordant increase in RhoA activity. The balance between these pathways depends on p120ctn levels and determines the amount of E-cad endocytosis. Therefore, we suggest that p120ctn co-ordinates cell adhesion with endocytosis and the actin cytoskeleton to dictate cellular behaviour and tissue tension.

Previous work established a cap model whereby p120ctn prevents the internalization of E-cad in mammalian cells [32,93–95]. This function is ascribed to two motifs in mammalian E-cad (LL and DEE), which are absent in *Drosophila* E-cad [32]. This explains why *Drosophila* E-cad is not completely internalised in *p120ctn* mutants [33]. Here, we showed that p120ctn overexpression inhibits E-cad endocytosis. Superficially, this is similar to mammalian cells, in which stabilization results from p120ctn outcompeting access to the E-cad LL and DEE motifs [96]. However, our findings represent a distinct phenomenon resulting from the perturbed balance of downstream signalling. Indeed, it is more similar to the mechanism of E-cad internalisation in mammalian cells, which does not require p120ctn dissociation from E-cad, but instead relies on recruitment of clathrin adaptor AP2 through direct interaction between p120ctn and Numb [25]. An outstanding question is whether this mechanism in mammalian cells also involves regulation of actin remodelling by p120ctn.

Both RhoA and Arf1 regulate actin remodelling and endocytosis, but their functions are fundamentally different. RhoA has been one of the most well studied elements of the regulated remodelling of the actin cytoskeleton [97]. Its distribution and recruitment of various components lead to increasing cortical tension via the phosphorylation of non-muscle Myosin II. In turn, increased cortical tension counteracts membrane bending which is required for vesicle formation and endocytosis [98,99]. In mammalian cells, p120ctn is a well-established regulator of RhoA, and in most cases either directly or indirectly inhibits it, which plays an important role in regulation of epithelial cell shape [26,100]. However, it was unclear if this p120ctn function is present in *Drosophila*: p120ctn physically binds RhoA, but is not required for RhoA localization and does not genetically interact with it [50,51]. Here, we demonstrated that Rho activity positively correlates with p120ctn levels at the membrane. An interesting question is whether *Drosophila* p120ctn can also inhibit RhoA or can localize its activity in other tissues or developmental contexts, and whether activation of RhoA is an ancestral function. If so, this would imply that RhoA inhibition is a more novel acquisition in chordates which arose during the diversification of the p120ctn family [10] necessitated by the increased complexity and variety of mammalian tissues and by the longevity of mammalian organisms. Further we have shown that the functional outcome of this p120ctn-RhoA interaction is to stabilize E-cad. Elevated p120ctn and RhoA activity impairs the formation of vesicles to internalize E-cad, and therefore increases the duration E-cad spends at the membrane, shifting the set point of steady-state E-cad membrane levels.

In contrast, Arf1 promotes E-cad endocytosis by both recruiting clathrin adaptor proteins and facilitating the remodelling of actin required for vesicle formation and abscission. In this study we showed for the first time a role for Arf1 in the clathrin-mediated trafficking of E-cad from the membrane. Prior to our work the Arf1 recruitment of clathrin had only been shown to occur at the Golgi by recruiting the Adaptor 1 protein (AP1) [84]. The function of Arf1 at the plasma membrane had been descried in dynamin-independent endocytosis, which was presumed to be clathrin-independent [101]. However, the involvement of Arf1 in multiple endocytic pathways has not been fully explored. We have shown that Arf1’s capacity to recruit clathrin is exploited by p120ctn to facilitate the endocytosis of E-cad. Whether this requires AP2, a well-documented membrane resident clathrin adaptor, has yet to be determined. Our findings provide a mechanistic insight into the pro-endocytic activity of p120ctn which has only recently come to light [25,39] and elaborates the number of known p120ctn interactors. The activities of Arf1 and RhoA are antagonistic, which was also seen in previous studies on the cellularization of the early embryo [44].

We propose that by balancing the RhoA and Arf1 pathways p120ctn determines the precise amount of E-cad endocytosis. Further, using laser ablation we demonstrated that p120ctn directly modulates tension, providing a broader view for the function of p120ctn on the tissue-level. Considering that all of the components of this regulatory network are expressed in all epithelia across evolution, we speculate that this system is likely to be more broadly applicable in development and a general feature of cell biology.

## Materials and Methods

### Fly stocks and genetics

Flies were raised on standard medium. The GAL4/UAS system [102] was used for all the specific spatial and temporal expression of transgenic and RNAi experiments. The GAL4 expressional driver used for all experiments was *engrailed∷GAL4* (*en*∷GAL4, Bloomington number 30564). The following fly stocks were used in this study (Bloomington number included where applicable): E-cad-GFP (*shg∷E-cadherin-GFP*) (60584), Shg-Cherry (*shg∷mCherry*, 59014), *UAS*∷CLC-GFP (7109), *UAS*∷Arf1-GFP (gift from T.Harris), Zipper-YFP (*Myosin II-*YFP, Kyoto Stock Center 115082), *sqh*∷Rok^K116A^-Venus (gift from J.Zallen), *UAS*∷Arf1-T31N (DN) and *UAS∷*Arf1-Q71L (CA) [83], *UAS*∷Rho1-N19 (DN) (7328), and *tubulin*∷GAL80^TS^ (7017). The p120ctn mutant genotype (p120ctn^308^/ Δp120) used was derived from crossing two deficiency stocks: homozygously viable p120ctn^308^ females [33] with homozygously lethal Df(2R)M41A8/ CyO, *twi∷*Gal4, *UAS∷*GFP males (740). Thus, the p120ctn mutants examined lacked both maternal and zygotic contributions.

### Embryo collection and fixation

Embryos were collected at 25°C at 3-hour time intervals and allowed to develop at 18°C for 21 hours to reach the desired developmental stage, except for the acute induction experiments, where embryos were allowed to develop for 13 hours at 18°C and shifted to 29°C for 4 hours. Then embryos were dechorionated using 50% sodium hypochlorite (bleach, Invitrogen) in water for 4 minutes, and extensively washed with deionized water prior to fixation. Fixation was performed with a 1:1 solution of 4% formaldehyde (Sigma) in PBS (Phosphate Buffered Saline) and heptane (Sigma) for 20 minutes on an orbital shaker at room temperature. Embryos were then devitellinized in 1:1 solution of methanol and heptane for 20 sec with vigorous agitation. Following subsequent methanol washes the fixed embryo specimens were stored at −20°C in methanol until required.

### Embryo live imaging

Embryos were collected and dechorionated as described above. Once washed with deionized water embryos were transferred to apple juice agar segments upon microscope slide. Correct genotypes were selected under a fluorescent microscope (Leica) using a needle. Embryos were positioned and orientated in a row consisting of 6-10 embryos per genotype. Following this, embryos were transferred to pre-prepared microscope slides with Scotch tape and embedded in Halocarbon oil 27 (Sigma). Embryos were left to aerate for 10 minutes prior to covering with a cover slip and imaging.

For laser ablation, following orientation and positioning the embryos were transferred to a 60mm x 22mm coverslip which had been pre-prepared by applying 10 µl of Heptane glue along a strip in the middle of the coverslip orientated with the long axis. The coverslip was attached to a metal slide cassette (Zeiss), and the embryos were embedded in Halocarbon oil 27 before imaging.

### Molecular cloning

The p120ctn full length cDNA was obtained from Berkeley Drosophila Genome Project (BDGP), supplied in a pBSSK vector. This was sub-cloned into a (pUAS-k10.attB) plasmid using standard restriction digestion with NotI and BamHI (New England Biolabs) followed by ligation with T4 DNA ligase (New England Biolabs) and transformation into DH5a competent E.coli cells (Thermo Fisher Scientific). Prior to injection plasmids were test digested and sequenced (Core Genomic Facility, University of Sheffield). Plasmids were prepared for injection using standard miniprep extraction (Qiagen) and submitted for injection (Microinjection service, Department of Genetics, University of Cambridge) into the attP-86Fb stock (Bloomington stock 24749). Successful incorporation of the transgene was determined by screening for (*w*^+^) in the F1 progeny.

### Immunostaining

The embryos were washed three times in 1 ml of PBST (PBS with 0.05% Triton) with gentle rocking. Blocking of the embryos prior to staining was done in 300 µl of a 1% NGS (Normal Goat Serum) in PBST for 1 hour at room temperature with gentle rocking. For staining the blocking solution was removed, 300 µl of the primary antibody: either 1:100 dilution of a rat anti-E-cad (DCAD2, DSHB) or 1:10 of a mouse anti-engrailed (4D9, DSHB) in fresh blocking solution was added and the embryos were incubated overnight at 4°C with orbital rotation. Then, embryos were washed three times with 1 ml of PBST. A 300 µl 1:300 dilution of the secondary antibody (goat Cy3- or Cy5-conjugated anti-rat-IgG, Invitrogen) was added, and the embryos incubated either overnight at 4°C with orbital rotation or for 2 hours at room temperature with gentle rocking. Then embryos were washed three time with PBST, following which they were incubated with 50-70 µl of Vectashield (Vector Laboratories) and allowed to equilibrate for a period of 2 hours before being mounted on microscope slides (Thermo).

### Microscopy, data acquisition and FRAP

All experiments except for laser ablation were performed using an up-right Olympus FV1000 confocal microscope with a 60x/1.40 NA oil immersion objective. All measurements were made on dorsolateral epidermal cells of embryos, which were near or just after completion of dorsal closure, corresponding to the end of Stage 15 of embryogenesis. For fixed samples 16-bit images were taken at a magnification of 0.051µm/pixel (1024×1024 pixel XY-image) with a pixel dwell of 4µm/pixel. For each embryo, a Z-axis sectional stack through the plane of the AJs was taken, which consisted of six sections with a 0.38 µm intersectional spacing. The images were saved in the Olympus binary image format for further processing.

For E-cad FRAP (adapted from Bulgakova et al., 2013) 16-bit images were taken at a magnification of 0.093 µm/pixel (320×320 pixel XY-image). In each embryo, several circular regions of 1 µm radius were photobleached at either DV or AP junctions resulting in one bleach event per cell. Photobleaching was performed with 8 scans at 2 µs/pixel at 50-70% 488 nm laser power, resulting in the reduction of E-cad-GFP signal by 60–80%. A stack of 6 z-sections spaced by 0.38 µm was imaged just before photobleaching, and immediately after photobleaching, and then at 20 s intervals, for a total of 15 minutes.

As rate of endocytosis depends on external factors, such as temperature [103], controls were examined in parallel with experimental conditions in all experiments with CLC-GFP. This dependence might also account for slight variation between datasets, e.g. compare Fig. 2A and 2G. For CLC-GFP FRAP, 16-bit images were taken at a magnification of 0.051µm/pixel (256×256 pixel XY-image). In each embryo a single plane was selected in centre of the AJ band using Shg-Cherry for positioning. An area encompassing a transverse region orthogonal to the axis of the engrailed expressing cells was selected (140×60 pixels) was photobleached with 1 scan at 2 µm/pixel using 100% 488nm laser power resulting in reduction of CLC-GFP signal by 70-80%. Images were taken using continuous acquisition at a frame rate of 2 sec^-1^. Prior to bleaching a sequence of 10 images was taken, and a total of 400 frames corresponding to 3.5 minutes were taken.

### Data processing and statistical analysis

#### Membrane intensity

Images were processed in Fiji (https://fiji.sc) by generating average intensity projections of the channel required for quantification. Masks were created by processing background-subtracted maximum intensity projections using the Tissue Analyzer plugin in Fiji [104]. Quantification of the membrane intensity at the AP and DV borders was done as described previously using a custom-built Matlab script [39] found at (https://github.com/nbul/Intensity). In the case of quantification of Arf1-GFP membrane intensity, due to high Arf1-GFP presence inside cells both in Golgi and cytoplasm, the mean intensity of embryonic areas not expressing the GFP-tagged transgene were used for background subtraction. Statistical analysis was performed in Graphpad Prism (https://www.graphpad.com/scientific-software/prism/). First, the data was cleaned using ROUT detection of outliers in Prism followed by testing for normal distribution (D’Agostino & Pearson normality test). Then, the significance for parametric data was tested by either a two-way ANOVA or two-tailed t-test with Welch’s correction.

#### E-cad FRAP

images were processed by using the grouped Z-projector plugin in Fiji to generate average intensity projections for each time-point. Following this the bleached ROI, control ROI and background intensity were manual measured for each time point. This data was processed in Microsoft Excel. First the intensity of the bleached ROI at each time point was background subtracted and normalized as following: *I*_*n*_ = (*F*_*n*_ - *BG*_*n*_)/(*FC*_*n*_ - *BG*_*n*_), where *F*_*n*_ – intensity of the bleached ROI at the time point *n*, *FC*_*n*_ – intensity of the control unbleached ROI of the same size at the plasma membrane at the time point *n*, and *BG*_*n*_ – background intensity, measured with the same size ROI in cytoplasm at the time point *n*. Than the relative recovery at each time point was calculated using the following formula: *R*_*n*_ = (*I*_*n*_ - *I*_1_)/(*I*_0_ - *I*_1_), where *I*_*n*_, *I*_1_ and *I*_0_ are normalized intensities of bleached ROI and time point *n*, immediately after photobleaching, and before photobleaching respectively.

These values were input to Prism and nonlinear regression analysis was performed to test for best fit model and if recoveries were significantly different between cell borders or genotypes. The recovery was fit to either single exponential model in a form of 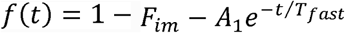, or to bi-exponential model in a form of 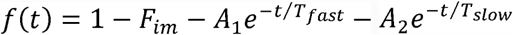, where *F*_*im*_ is a size of the immobile fraction, *T*_*fast*_ and *T*_*slow*_ are the half times, and *A*_1_ and *A*_2_ are amplitudes of the fast and slow components of the recovery. An F-test was used to choose the model and compare datasets.

#### CLC-GFP FRAP

measurements of all intensities, i.e. the bleached ROI, control ROI and the background, and normalization were done using a custom-build Matlab script (http://github.com/nbul/FRAP) using the same algorithm as described for E-cad FRAP. Curve fitting and statistical analysis was performed in Graphpad Prism using a nonlinear regression analysis as described for E-cad FRAP.

#### CLC-GFP puncta

Images were analysed using a custom script in MATLAB described in [64]. This was modified for unpaired data by calculating a threshold value for puncta detection using a mean calculated from all of the control images and applying this threshold to the experimental images. Statistical analysis of the recovery was performed in Graphpad Prism using nonlinear regression analysis.

#### Total membrane intensity

the masks generated using the Tissue analyser plugin were processed using a custom-built Matlab script (https://github.com/nbul/Intensity/tree/master/TissueAnalyzer). In short, the outlines of the cells were determined from the binary masks, and the length of outline was used as a measure of cell perimeter. Then, the outlines of individual cells were dilated using a YxY structuring element, so that the resulting mask covered all visible E-cad signal on the borders of the cell, and the mean pixel intensity of the mask was measured. The total protein amount was calculated as a product of cell perimeter by mean pixel intensity of the mask. Statistical analysis was performed in Graphpad Prism: using a two-tailed t-test with Welch’s correction.

### Laser Ablation

Nanoablation of single junctions was performed to provide a measure of junctional tension. Embryos were imaged on a Zeiss LSM 880 microscope with an Airyscan detector, an 8-bit image at 0.053 µm/pixel (512×512 pixel XY-Image) resolution with a 63x objective (NA 1.4) at 5x zoom and 2x averaging was used. An illumination wavelength of 488 nm and 0.5% laser power were used. Images were captured with a 0.5 µm z-spacing. Narrow rectangular ROIs were drawn across the centre of single junctions and this region was ablated using a pulsed TiSa laser (Chameleon), tuned to 760 nm at 45% power. Embryos were imaged continuously in a z-stack consisting of 3 z-slices. The initial recoil rate of vertices at the ends of ablated junctions was quantified by measuring the change in distance between the vertices and dividing by the initial time step. Statistical analysis was performed in Graphpad Prism: using a two-tailed t-test with Welch’s correction.

## Acknowledgments

The authors first wish to thank Rob Tetley and Yanlan Mao for assistance with laser ablation experiments. The authors also with to acknowledge the advice and guidance of Professor David Strutt, University of Sheffield, and his lab. The authors thank to the technical staff of the Wolfson Light Microscopy Facility (LMF) and the Fly Facility, the University of Sheffield, without whom this work would not be possible. This work was supported by grant BB/P007503/1 from the UK Biotechnology and Biological Sciences Research Council.

## Author Contribution

J.G and N.B. designed and performed experiments and wrote the manuscript.

## Supplementary Information

**Table S1. Numerical values for each experiment presented in paper.**

**Figure S1. Shg-GFP is less dynamic at both cell borders in the p120ctn mutant.**

**A-B.** Recovery of fluorescent intensity during the course of a FRAP experiment for the control (**A**) and p120ctn mutant embryos (**B**), measured at both the AP and DV cell borders. Average recovery curves (mean ± s.e.m.) and the best-fit curves (solid lines) are shown. All best-fit and membrane intensity data are in Table S1.

**Figure S2. Rho^CA^ induces substantial changes in cellular morphology and a decrease in Shg-GFP at the membrane.**

**A-B**. Apical view of the dorsolateral epidermis of control embryos (expressing CD8-m-Cherry transgene, **A-A’**, magenta in **A**), and embryos expressing RhoA^CA^ transgene with *en*∷Gal4 (B-B’, visualized with CD8-m-Cherry, magenta in **A**). Lower magnification images presented with white square representing region are shown in main figure (see Fig. 4). The cell membranes marked by E-cad-GFP are shown in green (**A, B**) and black (**A’,B’**). **C.** Histograms of the cell area, comparing engrailed regions of control and RhoA^CA^ expressing cells: individual distributions (**C**) and overlaid frequency distributions (**C’**). **D.** Histograms of the elongation of the cells, shown as individual (**D**) and overlaid (**D’**). **E.** Histograms of the distribution of cell neighbours, both as individual distributions (**E**) and overlaid curves (**E**’). **F.** Quantification of the amount of Shg-GFP at the cell membrane, calculating total membrane without angle dependence (division into AP and DV borders). Statistical analysis is a two-tailed students t-test with Welches correction *, P < 0.05; **, P < 0.001; ***, P < 0.0001. Scale bar is 10 µm.

**Figure S3. Arf1-GFP cytoplasmic puncta are representative of the Golgi apparatus.**

**A-E.** Apical view of the dorsolateral epidermis of *UAS∷*Arf1-GFP expressing control embryo with cells borders are marked by antibody staining of E-cad (black in **A**, white in **E**), Golgi marked by immunostaining of Trans-Golgi (black in **B**, magenta in D and E), and Arf1-GFP localization in the engrailed expressing cells (black in **C,** green in **D** and **E)**. The same region is shown in main text (see Fig. 5). Scale bar is 10 µm.

**Figure S4. Arf1^DN^ induces cell morphology changes leading to cell death.**

**A-B.** Apical view of the dorsolateral epidermis of Arf1^DN^ transgene expressing embryos, co-expressing *UAS∷*CD8-cherry in the engrailed stripes to mark the cells (magenta in **A**). The cell borders are marked by Shg-GFP (green in A, black in B). Scale bar is 10 µm. **C.** Histograms of the frequency distributions of the cell area between the Arf1^DN^ expressing cells and the adjacent internal control cells, shown as both the individual distributions (**C**) and overlaid (**C’**). **D.** Histograms of the frequency distributions of the cell elongations between the Arf1^DN^ and the internal control cell. Individual and overlaid to display modal shift. **E.** Histograms of the frequency distributions of the cell neighbours shown as individual (**E**) and overlaid (**E**’). Horizontal line represents 50% threshold for each cumulative distribution (**E’**). **F.** Histograms showing the cumulative distributions of the cell area showing the individual cumulative distributions (**F**) and the overlaid curves (**F’**).

## References

1. Wickström SA, Niessen CM,Niessen CM (2018) Cell adhesion and mechanics as drivers of tissue organization and differentiation: local cues for large scale organization. Curr Opin Cell Biol 54: 89–97.

2. van Roy F, Berx G (2008) The cell-cell adhesion molecule E-cadherin. Cell Mol Life Sci CMLS 65: 3756–3788.

3. Takeichi M (1977) Functional correlation between cell adhesive properties and some cell surface proteins. J Cell Biol 75: 464–474.

4. Ozawa M, Ringwald M, Kemler R (1990) Uvomorulin-catenin complex formation is regulated by a specific domain in the cytoplasmic region of the cell adhesion molecule. Proc Natl Acad Sci U S A 87: 4246–4250.

5. Shapiro L, Weis WI,Weis WI (2009) Structure and biochemistry of cadherins and catenins. Cold Spring Harb Perspect Biol 1: a003053.

6. Buckley CD,Buckley CD, Tan J, Anderson KL,Anderson KL, Hanein D, Volkmann N, Weis WI,Weis WI, Nelson WJ,Nelson WJ, Dunn AR,Dunn AR (2014) Cell adhesion. The minimal cadherin-catenin complex binds to actin filaments under force. Science 346: 1254211.

7. Daniel JM,Daniel JM, Reynolds AB,Reynolds AB (1995) The tyrosine kinase substrate p120cas binds directly to E-cadherin but not to the adenomatous polyposis coli protein or alpha-catenin. Mol Cell Biol 15: 4819–4824.

8. Shibamoto S, Hayakawa M, Takeuchi K, Hori T, Miyazawa K, Kitamura N, Johnson KR,Johnson KR, Wheelock MJ,Wheelock MJ, Matsuyoshi N, Takeichi M (1995) Association of p120, a tyrosine kinase substrate, with E-cadherin/catenin complexes. J Cell Biol 128: 949–957.

9. Yap AS,Yap AS, Niessen CM,Niessen CM, Gumbiner BM,Gumbiner BM (1998) The juxtamembrane region of the cadherin cytoplasmic tail supports lateral clustering, adhesive strengthening, and interaction with p120ctn. J Cell Biol 141: 779–789.

10. Carnahan RH,Carnahan RH, Rokas A, Gaucher EA,Gaucher EA, Reynolds AB,Reynolds AB (2010) The Molecular Evolution of the p120-Catenin Subfamily and Its Functional Associations. PLOS ONE 5: e15747.

11. Gul IS,Gul IS, Hulpiau P, Saeys Y, van Roy F (2017) Evolution and diversity of cadherins and catenins. Exp Cell Res 358: 3–9.

12. Hatzfeld M (2005) The p120 family of cell adhesion molecules. Eur J Cell Biol 84: 205–214.

13. Riethmacher D, Brinkmann V, Birchmeier C (1995) A targeted mutation in the mouse E-cadherin gene results in defective preimplantation development. Proc Natl Acad Sci 92: 855–859.

14. Shimizu T, Yabe T, Muraoka O, Yonemura S, Aramaki S, Hatta K, Bae Y-K, Nojima H, Hibi M (2005) E-cadherin is required for gastrulation cell movements in zebrafish. Mech Dev 122: 747–763.

15. Elisha Y, Kalchenko V, Kuznetsov Y, Geiger B (2018) Dual role of E-cadherin in the regulation of invasive collective migration of mammary carcinoma cells. Sci Rep 8: 4986.

16. Petrova YI,Petrova YI, Schecterson L, Gumbiner BM,Gumbiner BM (2016) Roles for E-cadherin cell surface regulation in cancer. Mol Biol Cell 27: 3233–3244.

17. Wells A, Yates C, Shepard CR,Shepard CR (2008) E-cadherin as an indicator of mesenchymal to epithelial reverting transitions during the metastatic seeding of disseminated carcinomas. Clin Exp Metastasis 25: 621–628.

18. Levayer R, Pelissier-Monier A, Lecuit T (2011) Spatial regulation of Dia and Myosin-II by RhoGEF2 controls initiation of E-cadherin endocytosis during epithelial morphogenesis. Nat Cell Biol 13: 529–540.

19. Troyanovsky RB,Troyanovsky RB, Sokolov EP,Sokolov EP, Troyanovsky SM,Troyanovsky SM, Nusrat A (2006) Endocytosis of Cadherin from Intracellular Junctions Is the Driving Force for Cadherin Adhesive Dimer Disassembly. Mol Biol Cell 17: 3484–3493.

20. Cadwell CM,Cadwell CM, Su W, Kowalczyk AP,Kowalczyk AP (2016) Cadherin Tales: Regulation of Cadherin Function by Endocytic Membrane Trafficking. Traffic Cph Den.

21. Garrett JP,Garrett JP, Lowery AM,Lowery AM, Adam AP,Adam AP, Kowalczyk AP,Kowalczyk AP, Vincent PA,Vincent PA (2017) Regulation of endothelial barrier function by p120-catenin·VE-cadherin interaction. Mol Biol Cell 28: 85–97.

22. Ireton RC,Ireton RC, Davis MA,Davis MA, van Hengel J, Mariner DJ,Mariner DJ, Barnes K, Thoreson MA,Thoreson MA, Anastasiadis PZ,Anastasiadis PZ, Matrisian L, Bundy LM,Bundy LM, Sealy L, et al. (2002) A novel role for p120 catenin in E-cadherin function. J Cell Biol 159: 465–476.

23. Oas RG,Oas RG, Nanes BA,Nanes BA, Esimai CC,Esimai CC, Vincent PA,Vincent PA, García AJ, Kowalczyk AP,Kowalczyk AP (2013) p120-catenin and β-catenin differentially regulate cadherin adhesive function. Mol Biol Cell 24: 704–714.

24. Reynolds AB,Reynolds AB (2007) p120-catenin: Past and present. Biochim Biophys Acta BBA - Mol Cell Res 1773: 2–7.

25. Sato K, Watanabe T, Wang S, Kakeno M, Matsuzawa K, Matsui T, Yokoi K, Murase K, Sugiyama I, Ozawa M, et al. (2011) Numb controls E-cadherin endocytosis through p120 catenin with aPKC. Mol Biol Cell 22: 3103–3119.

26. Yu HH,Yu HH, Dohn MR,Dohn MR, Markham NO,Markham NO, Coffey RJ,Coffey RJ, Reynolds AB,Reynolds AB (2016) p120-catenin controls contractility along the vertical axis of epithelial lateral membranes. J Cell Sci 129: 80–94.

27. Davis MA,Davis MA, Ireton RC,Ireton RC, Reynolds AB,Reynolds AB (2003) A core function for p120-catenin in cadherin turnover. J Cell Biol 163: 525–534.

28. Ishiyama N, Lee S-H, Liu S, Li G-Y, Smith MJ,Smith MJ, Reichardt LF,Reichardt LF, Ikura M (2010) Dynamic and Static Interactions between p120 Catenin and E-Cadherin Regulate the Stability of Cell-Cell Adhesion. Cell 141: 117–128.

29. Gold JS,Gold JS, Reynolds AB,Reynolds AB, Rimm DL,Rimm DL (1998) Loss of p120ctn in human colorectal cancer predicts metastasis and poor survival. Cancer Lett 132: 193–201.

30. Shibata T, Kokubu A, Sekine S, Kanai Y, Hirohashi S (2004) Cytoplasmic p120ctn regulates the invasive phenotypes of E-cadherin-deficient breast cancer. Am J Pathol 164: 2269–2278.

31. van de Ven RAH, Tenhagen M, Meuleman W, van Riel JJG, Schackmann RCJ, Derksen PWB (2015) Nuclear p120-catenin regulates the anoikis resistance of mouse lobular breast cancer cells through Kaiso-dependent Wnt11 expression. Dis Model Mech 8: 373–384.

32. Nanes BA,Nanes BA, Chiasson-MacKenzie C, Lowery AM,Lowery AM, Ishiyama N, Faundez V, Ikura M, Vincent PA,Vincent PA, Kowalczyk AP,Kowalczyk AP (2012) p120-catenin binding masks an endocytic signal conserved in classical cadherins. J Cell Biol 199: 365–380.

33. Myster SH,Myster SH, Cavallo R, Anderson CT,Anderson CT, Fox DT,Fox DT, Peifer M (2003) Drosophila p120catenin plays a supporting role in cell adhesion but is not an essential adherens junction component. J Cell Biol 160: 433–449.

34. Pacquelet A, Lin L, Rorth P (2003) Binding site for p120/delta-catenin is not required for Drosophila E-cadherin function in vivo. J Cell Biol 160: 313–319.

35. Pettitt J, Cox EA,Cox EA, Broadbent ID,Broadbent ID, Flett A, Hardin J (2003) The Caenorhabditis elegans p120 catenin homologue, JAC-1, modulates cadherin-catenin function during epidermal morphogenesis. J Cell Biol 162: 15–22.

36. Israely I, Costa RM,Costa RM, Xie CW,Xie CW, Silva AJ,Silva AJ, Kosik KS,Kosik KS, Liu X (2004) Deletion of the Neuron-Specific Protein Delta-Catenin Leads to Severe Cognitive and Synaptic Dysfunction. Curr Biol 14: 1657–1663.

37. Ho C, Zhou J, Medina M, Goto T, Jacobson M, Bhide PG,Bhide PG, Kosik KS,Kosik KS (2000) δ-catenin is a nervous system-specific adherens junction protein which undergoes dynamic relocalization during development. J Comp Neurol 420: 261–276.

38. Iyer KV,Iyer KV, Piscitello-Gómez R, Jülicher F, Eaton S (2018) Mechanosensitive binding of p120-Catenin at cell junctions regulates E-Cadherin turnover and epithelial viscoelasticity. bioRxiv 357186.

39. Bulgakova NA,Bulgakova NA, Brown NH,Brown NH (2016) Drosophila p120-catenin is crucial for endocytosis of the dynamic E-cadherin-Bazooka complex. J Cell Sci 129: 477–482.

40. Baum B, Georgiou M (2011) Dynamics of adherens junctions in epithelial establishment, maintenance, and remodeling. J Cell Biol 192: 907–917.

41. Cavey M, Lecuit T (2009) Molecular Bases of Cell–Cell Junctions Stability and Dynamics. Cold Spring Harb Perspect Biol 1:.

42. Smythe E, Ayscough KR,Ayscough KR (2006) Actin regulation in endocytosis. J Cell Sci 119: 4589–4598.

43. Derksen PWB, van de Ven RAH (2017) Shared mechanisms regulate spatiotemporal RhoA-dependent actomyosin contractility during adhesion and cell division. Small GTPases 1–9.

44. Lee DM,Lee DM, Harris TJC (2013) An Arf-GEF Regulates Antagonism between Endocytosis and the Cytoskeleton for Drosophila Blastoderm Development. Curr Biol 23: 2110–2120.

45. Anastasiadis PZ,Anastasiadis PZ, Moon SY,Moon SY, Thoreson MA,Thoreson MA, Mariner DJ,Mariner DJ, Crawford HC,Crawford HC, Zheng Y, Reynolds AB,Reynolds AB (2000) Inhibition of RhoA by p120 catenin. Nat Cell Biol 2: 637–644.

46. Lang RA,Lang RA, Herman K, Reynolds AB,Reynolds AB, Hildebrand JD,Hildebrand JD, Plageman TF,Plageman TF (2014) p120-catenin-dependent junctional recruitment of Shroom3 is required for apical constriction during lens pit morphogenesis. Dev Camb Engl 141: 3177–3187.

47. Taulet N, Comunale F, Favard C, Charrasse S, Bodin S, Gauthier-Rouvière C (2009) N-cadherin/p120 catenin association at cell-cell contacts occurs in cholesterol-rich membrane domains and is required for RhoA activation and myogenesis. J Biol Chem 284: 23137–23145.

48. Zebda N, Tian Y, Tian X, Gawlak G, Higginbotham K, Reynolds AB,Reynolds AB, Birukova AA,Birukova AA, Birukov KG,Birukov KG (2013) Interaction of p190RhoGAP with C-terminal domain of p120-catenin modulates endothelial cytoskeleton and permeability. J Biol Chem 288: 18290–18299.

49. Fox DT,Fox DT, Peifer M (2007) Cell Adhesion: Separation of p120’s Powers? Curr Biol 17: R24–R27.

50. Fox DT,Fox DT, Homem CCF, Myster SH,Myster SH, Wang F, Bain EE,Bain EE, Peifer M (2005) Rho1 regulates Drosophila adherens junctions independently of p120ctn. Development 132: 4819–4831.

51. Magie CR,Magie CR, Pinto-Santini D, Parkhurst SM,Parkhurst SM (2002) Rho1 interacts with p120ctn and α-catenin, and regulates cadherin-based adherens junction components in Drosophila. Development 129: 3771–3782.

52. Paterson AD,Paterson AD, Parton RG,Parton RG, Ferguson C, Stow JL,Stow JL, Yap AS,Yap AS (2003) Characterization of E-cadherin Endocytosis in Isolated MCF-7 and Chinese Hamster Ovary Cells THE INITIAL FATE OF UNBOUND E-CADHERIN. J Biol Chem 278: 21050–21057.

53. Donaldson JG,Donaldson JG, Jackson CL,Jackson CL (2011) ARF family G proteins and their regulators: roles in membrane transport, development and disease. Nat Rev Mol Cell Biol 12: 362–375.

54. McMahon HT,McMahon HT, Boucrot E (2011) Molecular mechanism and physiological functions of clathrin-mediated endocytosis. Nat Rev Mol Cell Biol 12: 517–533.

55. Humphreys D, Davidson AC,Davidson AC, Hume PJ,Hume PJ, Makin LE,Makin LE, Koronakis V (2013) Arf6 coordinates actin assembly through the WAVE complex, a mechanism usurped by Salmonella to invade host cells. Proc Natl Acad Sci 110: 16880–16885.

56. Padovani D, Folly-Klan M, Labarde A, Boulakirba S, Campanacci V, Franco M, Zeghouf M, Cherfils J (2014) EFA6 controls Arf1 and Arf6 activation through a negative feedback loop. Proc Natl Acad Sci U S A 111: 12378–12383.

57. Humphreys D, Liu T, Davidson AC,Davidson AC, Hume PJ,Hume PJ, Koronakis V (2012) The Drosophila Arf1 homologue Arf79F is essential for lamellipodium formation. J Cell Sci 125: 5630–5635.

58. Rodrigues FF,Rodrigues FF, Shao W, Harris TJC (2016) The Arf GAP Asap promotes Arf1 function at the Golgi for cleavage furrow biosynthesis in Drosophila. Mol Biol Cell 27: 3143–3155.

59. Shao W, Wu J, Chen J, Lee DM,Lee DM, Tishkina A, Harris TJC (2010) A modifier screen for Bazooka/PAR-3 interacting genes in the Drosophila embryo epithelium. PloS One 5: e9938.

60. Toret CP,Toret CP, D’Ambrosio MV, Vale RD,Vale RD, Simon MA,Simon MA, Nelson WJ,Nelson WJ (2014) A genome-wide screen identifies conserved protein hubs required for cadherin-mediated cell–cell adhesion. J Cell Biol 204: 265–279.

61. Adams CL,Adams CL, Nelson WJ,Nelson WJ, Smith SJ,Smith SJ (1996) Quantitative analysis of cadherin-catenin-actin reorganization during development of cell-cell adhesion. J Cell Biol 135: 1899–1911.

62. Tepass U, Hartenstein V (1994) The development of cellular junctions in the Drosophila embryo. Dev Biol 161: 563–596.

63. Coffman VC,Coffman VC, Wu J-Q (2012) Counting protein molecules using quantitative fluorescence microscopy. Trends Biochem Sci 37: 499–506.

64. Strutt H, Gamage J, Strutt D (2016) Robust Asymmetric Localization of Planar Polarity Proteins Is Associated with Organization into Signalosome-like Domains of Variable Stoichiometry. Cell Rep 17: 2660–2671.

65. Bulgakova NA,Bulgakova NA, Grigoriev I, Yap AS,Yap AS, Akhmanova A, Brown NH,Brown NH (2013) Dynamic microtubules produce an asymmetric E-cadherin–Bazooka complex to maintain segment boundaries. J Cell Biol 201: 887–901.

66. Lippincott-Schwartz J, Altan-Bonnet N, Patterson GH,Patterson GH (2003) Photobleaching and photoactivation: following protein dynamics in living cells. Nat Cell Biol Suppl: S7–14.

67. de Beco S, Gueudry C, Amblard F, Coscoy S (2009) Endocytosis is required for E-cadherin redistribution at mature adherens junctions. Proc Natl Acad Sci U S A 106: 7010–7015.

68. Xiao K, Allison DF,Allison DF, Buckley KM,Buckley KM, Kottke MD,Kottke MD, Vincent PA,Vincent PA, Faundez V, Kowalczyk AP,Kowalczyk AP (2003) Cellular levels of p120 catenin function as a set point for cadherin expression levels in microvascular endothelial cells. J Cell Biol 163: 535–545.

69. Pieters T, Goossens S, Haenebalcke L, Andries V, Stryjewska A, De Rycke R, Lemeire K, Hochepied T, Huylebroeck D, Berx G, et al. (2016) p120 Catenin-Mediated Stabilization of E-Cadherin Is Essential for Primitive Endoderm Specification. PLoS Genet 12: e1006243.

70. Chang HC,Chang HC, Newmyer SL,Newmyer SL, Hull MJ,Hull MJ, Ebersold M, Schmid SL,Schmid SL, Mellman I (2002) Hsc70 is required for endocytosis and clathrin function in Drosophila. J Cell Biol 159: 477–487.

71. Gaidarov I, Santini F, Warren RA,Warren RA, Keen JH,Keen JH (1999) Spatial control of coated-pit dynamics in living cells. Nat Cell Biol 1: 1–7.

72. Kochubey O, Majumdar A, Klingauf J (2006) Imaging Clathrin Dynamics in Drosophila melanogaster Hemocytes Reveals a Role for Actin in Vesicle Fission. Traffic 7: 1614–1627.

73. Hoffmann A, Dannhauser PN,Dannhauser PN, Groos S, Hinrichsen L, Curth U, Ungewickell EJ,Ungewickell EJ (2010) A comparison of GFP-tagged clathrin light chains with fluorochromated light chains in vivo and in vitro. Traffic Cph Den 11: 1129–1140.

74. Kaksonen M, Roux A (2018) Mechanisms of clathrin-mediated endocytosis. Nat Rev Mol Cell Biol 19: 313–326.

75. Leung T, Manser E, Tan L, Lim L (1995) A novel serine/threonine kinase binding the Ras-related RhoA GTPase which translocates the kinase to peripheral membranes. J Biol Chem 270: 29051–29054.

76. Amano M, Nakayama M, Kaibuchi K (2010) Rho-Kinase/ROCK: A Key Regulator of the Cytoskeleton and Cell Polarity. Cytoskelet Hoboken Nj 67: 545–554.

77. Schwayer C, Sikora M, Slováková J, Kardos R, Heisenberg C-P (2016) Actin Rings of Power. Dev Cell 37: 493–506.

78. Shewan AM,Shewan AM, Maddugoda M, Kraemer A, Stehbens SJ,Stehbens SJ, Verma S, Kovacs EM, Kovacs EM, Yap AS,Yap AS (2005) Myosin 2 Is a Key Rho Kinase Target Necessary for the Local Concentration of E-Cadherin at Cell–Cell Contacts. Mol Biol Cell 16: 4531–4542.

79. de Matos Simões S, Blankenship JT,Blankenship JT, Weitz O, Farrell DL,Farrell DL, Tamada M, Fernandez-Gonzalez R, Zallen JA,Zallen JA (2010) Rho-kinase directs Bazooka/Par-3 planar polarity during Drosophila axis elongation. Dev Cell 19: 377–388.

80. Pilauri V, Bewley M, Diep C, Hopper J (2005) Gal80 Dimerization and the Yeast GAL Gene Switch. Genetics 169: 1903–1914.

81. Jian X, Cavenagh M, Gruschus JM,Gruschus JM, Randazzo PA,Randazzo PA, Kahn RA,Kahn RA (2010) Modifications to the C-terminus of Arf1 alter cell functions and protein interactions. Traffic Cph Den 11: 732–742.

82. Ren X, Farias GG,Farias GG, Canagarajah BJ,Canagarajah BJ, Bonifacino JS,Bonifacino JS, Hurley JH,Hurley JH (2013) Structural Basis for Recruitment and Activation of the AP-1 Clathrin Adaptor Complex by Arf1. Cell 152: 755–767.

83. Wang Y, Zhang H, Shi M, Liou Y-C, Lu L, Yu F (2017) Sec71 functions as a GEF for the small GTPase Arf1 to govern dendrite pruning of Drosophila sensory neurons. Development 144: 1851–1862.

84. Carvajal-Gonzalez JM, Balmer S, Mendoza M, Dussert A, Collu G, Roman A-C, Weber U, Ciruna B, Mlodzik M (2015) The clathrin adaptor AP-1 complex and Arf1 regulate planar cell polarity in vivo. Nat Commun 6: 6751.

85. Luchsinger C, Aguilar M, Burgos PV,Burgos PV, Ehrenfeld P, Mardones GA,Mardones GA (2018) Functional disruption of the Golgi apparatus protein ARF1 sensitizes MDA-MB-231 breast cancer cells to the antitumor drugs Actinomycin D and Vinblastine through ERK and AKT signaling. PloS One 13: e0195401.

86. Schlienger S, Campbell S, Claing A (2014) ARF1 regulates the Rho/MLC pathway to control EGF-dependent breast cancer cell invasion. Mol Biol Cell 25: 17–29.

87. Myers KR,Myers KR, Casanova JE,Casanova JE (2008) Regulation of actin cytoskeleton dynamics by Arf-family GTPases. Trends Cell Biol 18: 184–192.

88. Boulant S, Kural C, Zeeh J-C, Ubelmann F, Kirchhausen T (2011) Actin dynamics counteract membrane tension during clathrin-mediated endocytosis. Nat Cell Biol 13: 1124–1131.

89. Mao Y, Tournier AL,Tournier AL, Hoppe A, Kester L, Thompson BJ,Thompson BJ, Tapon N (2013) Differential proliferation rates generate patterns of mechanical tension that orient tissue growth. EMBO J 32: 2790–2803.

90. Liang X, Michael M, Gomez GA,Gomez GA (2016) Measurement of Mechanical Tension at Cell-cell Junctions Using Two-photon Laser Ablation. Bio-Protoc **6**:.

91. Reynolds AB,Reynolds AB, Roczniak-Ferguson A (2004) Emerging roles for p120-catenin in cell adhesion and cancer. Oncogene 23: 7947–7956.

92. Priya R, Gomez GA,Gomez GA, Budnar S, Verma S, Cox HL,Cox HL, Hamilton NA,Hamilton NA, Yap AS,Yap AS (2015) Feedback regulation through myosin II confers robustness on RhoA signalling at E-cadherin junctions. Nat Cell Biol 17: 1282–1293.

93. Kowalczyk AP,Kowalczyk AP, Reynolds AB,Reynolds AB (2004) Protecting your tail: regulation of cadherin degradation by p120-catenin. Curr Opin Cell Biol 16: 522–527.

94. Su W, Kowalczyk AP,Kowalczyk AP (2016) The VE-cadherin cytoplasmic domain undergoes proteolytic processing during endocytosis. Mol Biol Cell.

95. Thoreson MA,Thoreson MA, Anastasiadis PZ,Anastasiadis PZ, Daniel JM,Daniel JM, Ireton RC,Ireton RC, Wheelock MJ,Wheelock MJ, Johnson KR,Johnson KR, Hummingbird DK,Hummingbird DK, Reynolds AB,Reynolds AB (2000) Selective Uncoupling of P120ctn from E-Cadherin Disrupts Strong Adhesion. J Cell Biol 148: 189–202.

96. Hartsock A, Nelson WJ,Nelson WJ (2012) Competitive Regulation of E-Cadherin JuxtaMembrane Domain Degradation by p120-Catenin Binding and Hakai-Mediated Ubiquitination. PLOS ONE 7: e37476.

97. Hodge RG,Hodge RG, Ridley AJ,Ridley AJ (2016) Regulating Rho GTPases and their regulators. Nat Rev Mol Cell Biol 17: 496–510.

98. Curtis Ide, Meldolesi J (2012) Cell surface dynamics – how Rho GTPases orchestrate the interplay between the plasma membrane and the cortical cytoskeleton. J Cell Sci 125: 4435–4444.

99. Jarsch IK,Jarsch IK, Daste F, Gallop JL,Gallop JL (2016) Membrane curvature in cell biology: An integration of molecular mechanisms. J Cell Biol 214: 375–387.

100. Menke A, Giehl K (2012) Regulation of adherens junctions by Rho GTPases and p120-catenin. Arch Biochem Biophys 524: 48–55.

101. Kumari S, Mayor S (2008) ARF1 is directly involved in dynamin-independent endocytosis. Nat Cell Biol 10: 30–41.

102. Brand AH,Brand AH, Perrimon N (1993) Targeted gene expression as a means of altering cell fates and generating dominant phenotypes. Dev Camb Engl 118: 401–415.

103. Weigel PH,Weigel PH, Oka JA,Oka JA (1981) Temperature dependence of endocytosis mediated by the asialoglycoprotein receptor in isolated rat hepatocytes. Evidence for two potentially rate-limiting steps. J Biol Chem 256: 2615–2617.

104. Aigouy B, Bivic AL,Bivic AL (2016) The PCP pathway regulates Baz planar distribution in epithelial cells. Sci Rep 6: srep33420.

