## Supplementary figures and images for "E-cadherin endocytosis is modulated by p120-catenin through the opposing actions of RhoA and Arf1"

### Supplementary figures

A

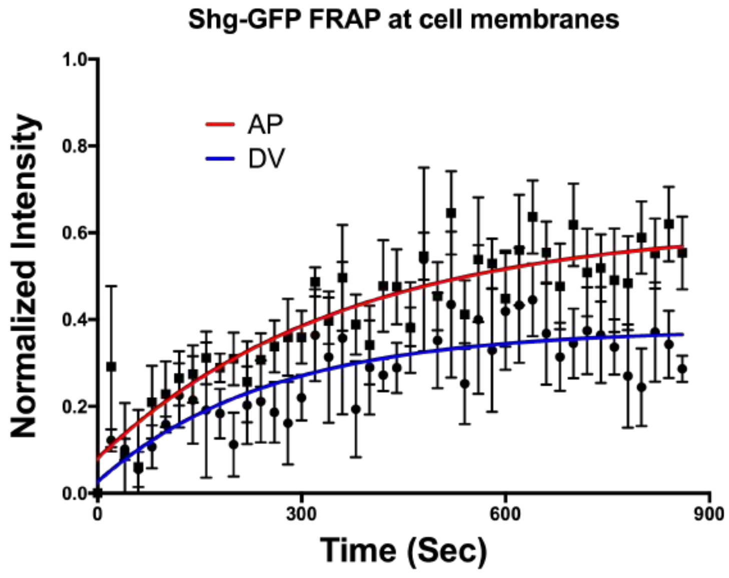


Control

B

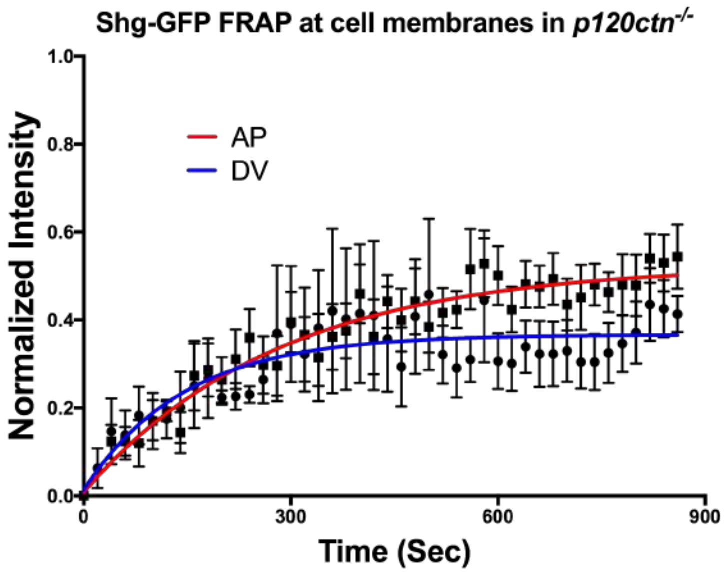


*p120ctn*$^{-/-}$

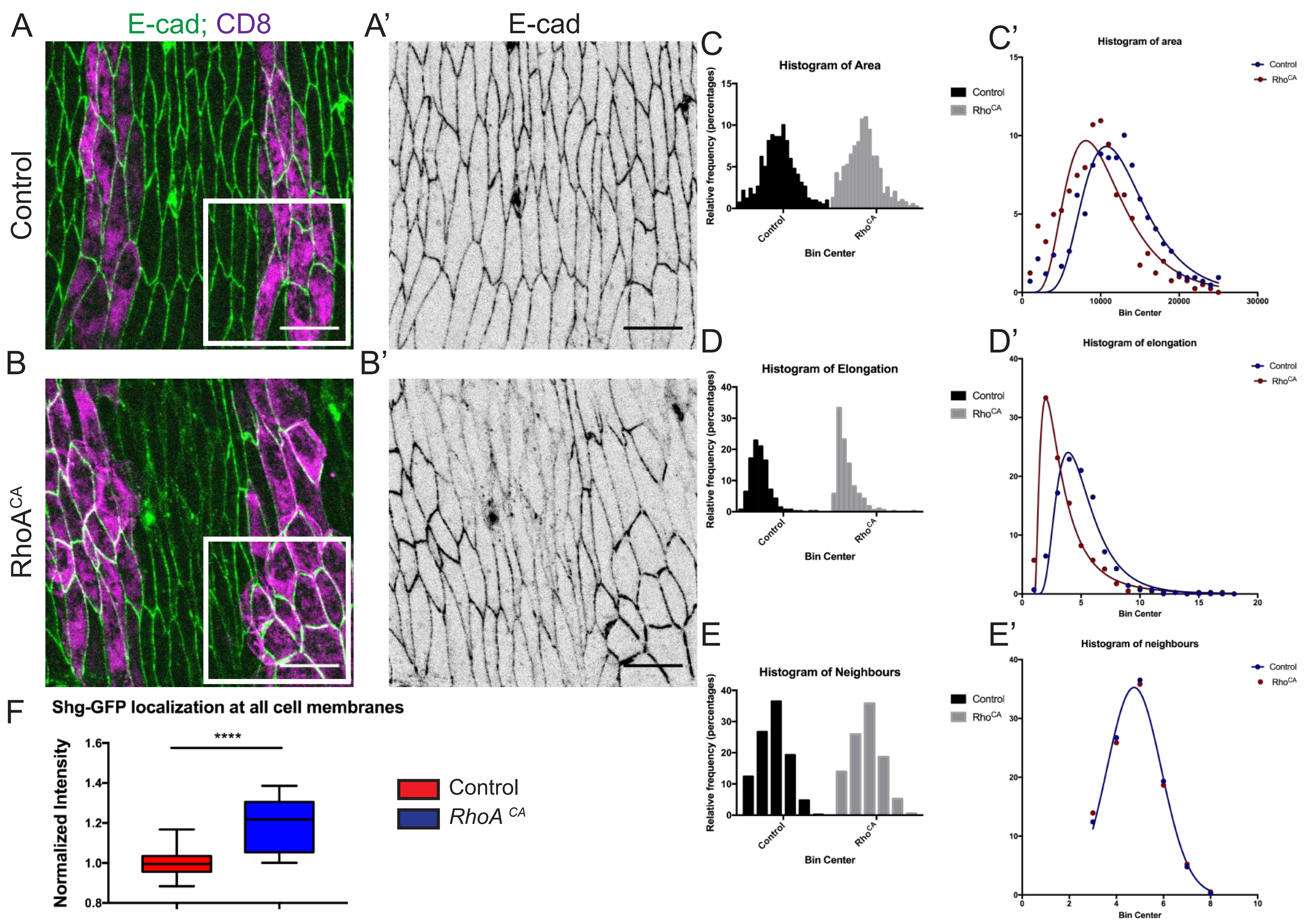

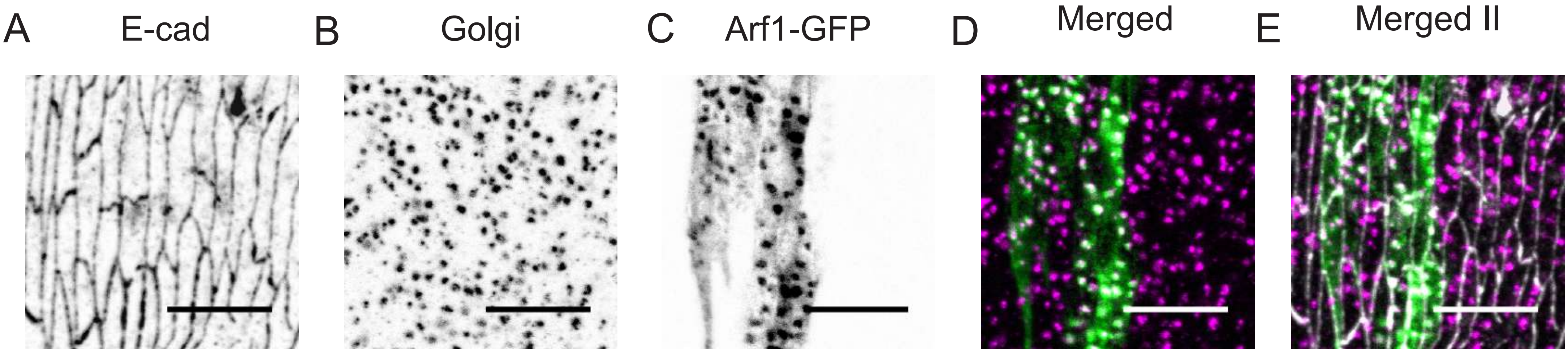

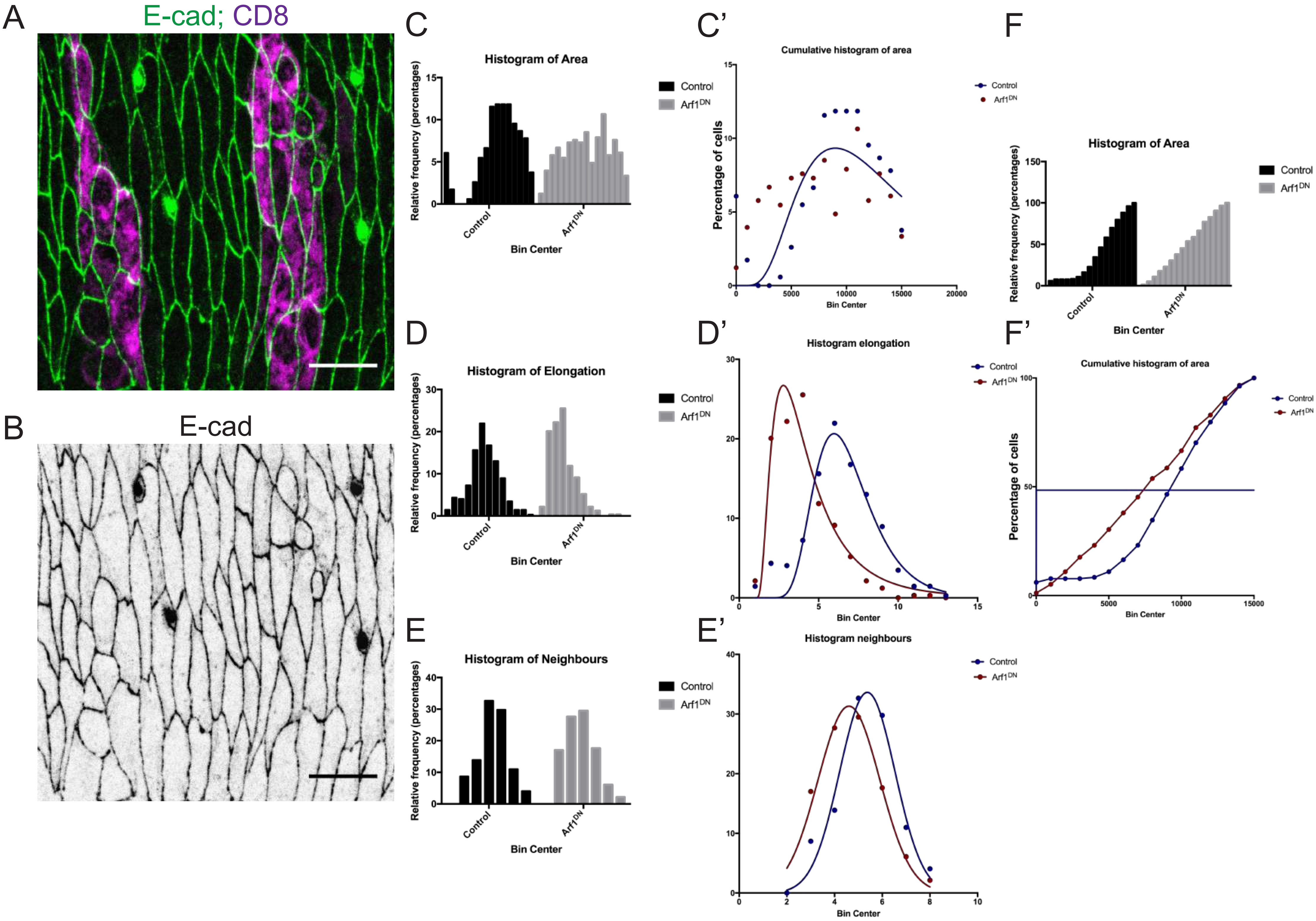
